# Rationally Designed Pooled CRISPRi-Seq Uncovers an Inhibitor of Bacterial Peptidyl-tRNA Hydrolase

**DOI:** 10.1101/2024.05.02.592284

**Authors:** A. S. M. Zisanur Rahman, Egor A. Syroegin, Julieta Novomisky Nechcoff, Archit Devarajan, Yury S. Polikanov, Silvia T. Cardona

## Abstract

Pooled knockdown libraries of essential genes are useful tools for elucidating the mechanisms of action of antibacterial compounds, a pivotal step in antibiotic discovery. However, achieving genomic coverage of antibacterial targets poses a challenge due to the uneven proliferation of knockdown mutants during pooled growth, leading to the unintended loss of important targets. To overcome this issue, we describe the construction of CIMPLE (<u>C</u>RISPR<u>i</u>-<u>m</u>ediated <u>p</u>ooled library of <u>e</u>ssential genes), a rationally designed pooled knockdown library built in a model antibiotic-resistant bacteria, *Burkholderia cenocepacia.* By analyzing growth parameters of clonal knockdown populations of an arrayed CRISPRi library, we predicted strain depletion levels during pooled growth and adjusted mutant relative abundance, approaching genomic coverage of antibacterial targets during antibiotic exposure. We first benchmarked CIMPLE by chemical-genetic profiling of known antibacterials, then applied it to an uncharacterized bacterial growth inhibitor from a new class. CRISPRi-Seq with CIMPLE, followed by biochemical validation, revealed that the novel compound targets the peptidyl-tRNA hydrolase (Pth). Overall, CIMPLE leverages the advantages of arrayed and pooled CRISPRi libraries to uncover unexplored targets for antibiotic action.

**Summary:** Bacterial mutant libraries in which antibiotic targets are downregulated are useful tools to functionally characterize novel antimicrobials. These libraries are used for chemical-genetic profiling as target-compound interactions can be inferred by differential fitness of mutants during pooled growth. Mutants that are functionally related to the antimicrobial mode of action are usually depleted from the pool upon exposure to the drug. Although powerful, this method can fail when the unequal proliferation of mutant strains before exposure causes mutants to fall below the detection level in the library pool. To address this issue, we constructed an arrayed essential gene mutant library (EGML) in the antibiotic-resistant bacterium *Burkholderia cenocepacia* using CRISPR interference (CRISPRi) and analyzed the growth parameters of individual mutant strains. We then modelled depletion levels during pooled growth and used the model to rationally design an optimized CRISPR interference-mediated pooled library of essential genes (CIMPLE). By adjusting the initial inoculum of the knockdown mutants, we achieved coverage of the bacterial essential genome with mutant sensitization. We exposed CIMPLE to a recently discovered antimicrobial of a novel class and discovered it inhibits the peptidyl-tRNA hydrolase, an essential bacterial enzyme. In summary, we demonstrate the utility of CIMPLE and CRISPRi-Seq to uncover the mechanism of action of novel antimicrobial compounds.

**Graphical abstract:** 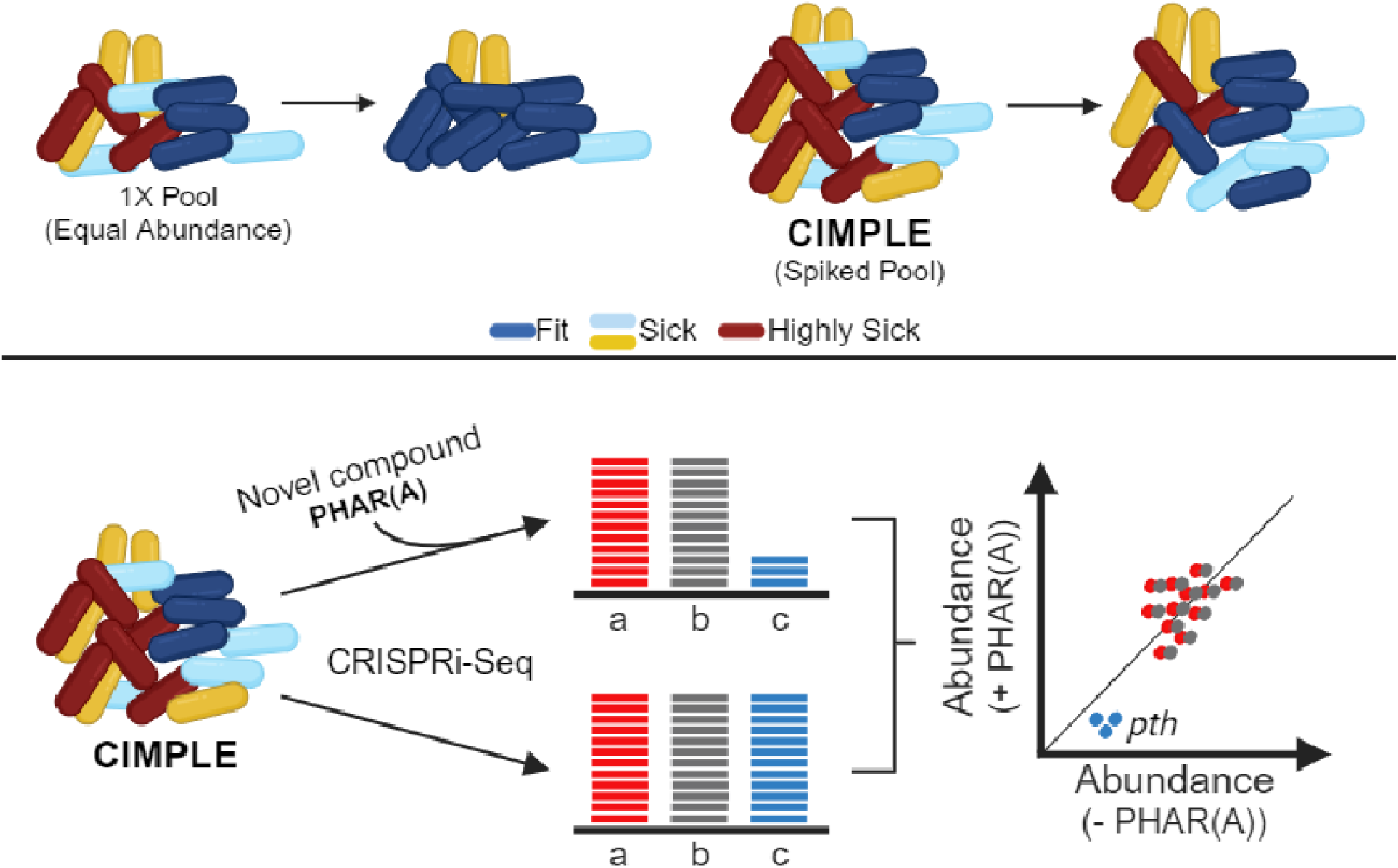

## Introduction

Antibiotic-resistant infections challenge public health and underscore the need for antibiotics of novel classes. In response, targeting the products of essential genes, which govern bacterial growth and viability, stands as one of the pivotal arms of early antibiotic drug discovery^1^. Essential gene mutant libraries (EGMLs) are useful tools to identify novel antimicrobials and determine their mechanism of action^2,3^. By downregulating an essential gene product, particular mutants are sensitized to compounds targeting that product or a related function. Consequently, the collective growth response of an EGML exposed to an antimicrobial can generate a chemical genetic profile of the drug, thus providing an initial characterization of its mechanism of action ^4,5^.

CRISPR interference (CRISPRi)^6^ is an ideal tool for creating EGMLs as it allows inducible repression of essential gene expression in bacteria^7^. CRISPRi utilizes a catalytically inactive Cas9 (dCas9) and a single guide RNA (sgRNA)^6^ to form a complex that binds to target DNA and sterically blocks transcription. Recognition of the target gene requires a short protospacer adjacent motif (PAM)^8–10^. CRISPRi can be applied to high throughput levels to create tunable^6,7^ and reversible^11^ CRISPRi mutant libraries. Several pooled^12–19^ and arrayed^7,20–22^ CRISPRi-based mutant libraries have been developed to facilitate the study of bacterial essential pathways^12,22–25^, identification of antibiotic susceptibility determinants^18,20,26–30^ and novel antimicrobial mechanisms of action^7^. Pooled libraries typically involve disrupting essential genes *en masse* and offer streamlined construction. However, these libraries are complex in number and risk disproportionate representation or even absence of essential gene mutants due to uneven growth. In contrast, arrayed libraries are smaller and have individual mutant strains representing the whole essential genome. However, profiling antimicrobials with arrayed CRISPRi libraries is costly as each mutant must be grown individually and exposed to the antimicrobials under investigation.

In this study, we describe the construction and use of CIMPLE (<u>C</u>RISPR<u>i</u>-<u>m</u>ediated <u>p</u>ooled library of <u>e</u>ssential genes), which combines the advantages of arrayed and pooled CRISPRi libraries. By quantifying the growth parameters of clonal mutant populations and modelling mutant depletion during pooled growth, we rationally manipulated the initial relative abundance of the mutants to create CIMPLE. Developed in the multiple antibiotic-resistant bacterium *Burkholderia cenocepacia* K56-2^31^, CIMPLE contains sensitized mutants covering the essential genome in a single pool. The CIMPLE platform, followed by CRISPRi-Seq, generated the chemical genetic profile of a novel growth inhibitor and signalled ribosome recycling as the essential function affected. Further biochemical characterization identified the peptidyl-tRNA hydrolase as the target of the novel compound. In summary, CIMPLE leverages the benefits of arrayed and pooled mutant libraries and demonstrates success in characterizing a novel antimicrobial mechanism of action.

## Results

### Construction and characterization of an arrayed essential gene knockdown mutant library (EGML) with CRISPRi technology

To construct the EGML in *B. cenocepacia* K56-2, we used our previously developed CRISPRi toolkit for *Burkholderia*^32^, in which a chromosomal copy of the *Streptococcus pyogenes* dCas9 (dCas9_Spy_) is expressed from a rhamnose-inducible promoter, and sgRNAs are constitutively expressed from a plasmid (**Fig. 1A**). We confirmed that the dCas9_Spy_ PAM preference allows more targetable regions in the *B. cenocepacia* K56-2 genome compared to other orthologous dCas9 proteins (**Supplementary Fig. 1**; **Supplementary Table 1**; **Supplementary Note 1**). To develop the arrayed CRISPRi library, we targeted the K56-2 essential genome comprising 535 genes^33,34^. Based on the predicted operons in the closely related *B. cenocepacia* J2315^35,36^ (**Supplementary Table 2)** and the polar effect of CRISPRi^6,7^, the number of targeted transcriptional units was 451.

**Fig. 1:**
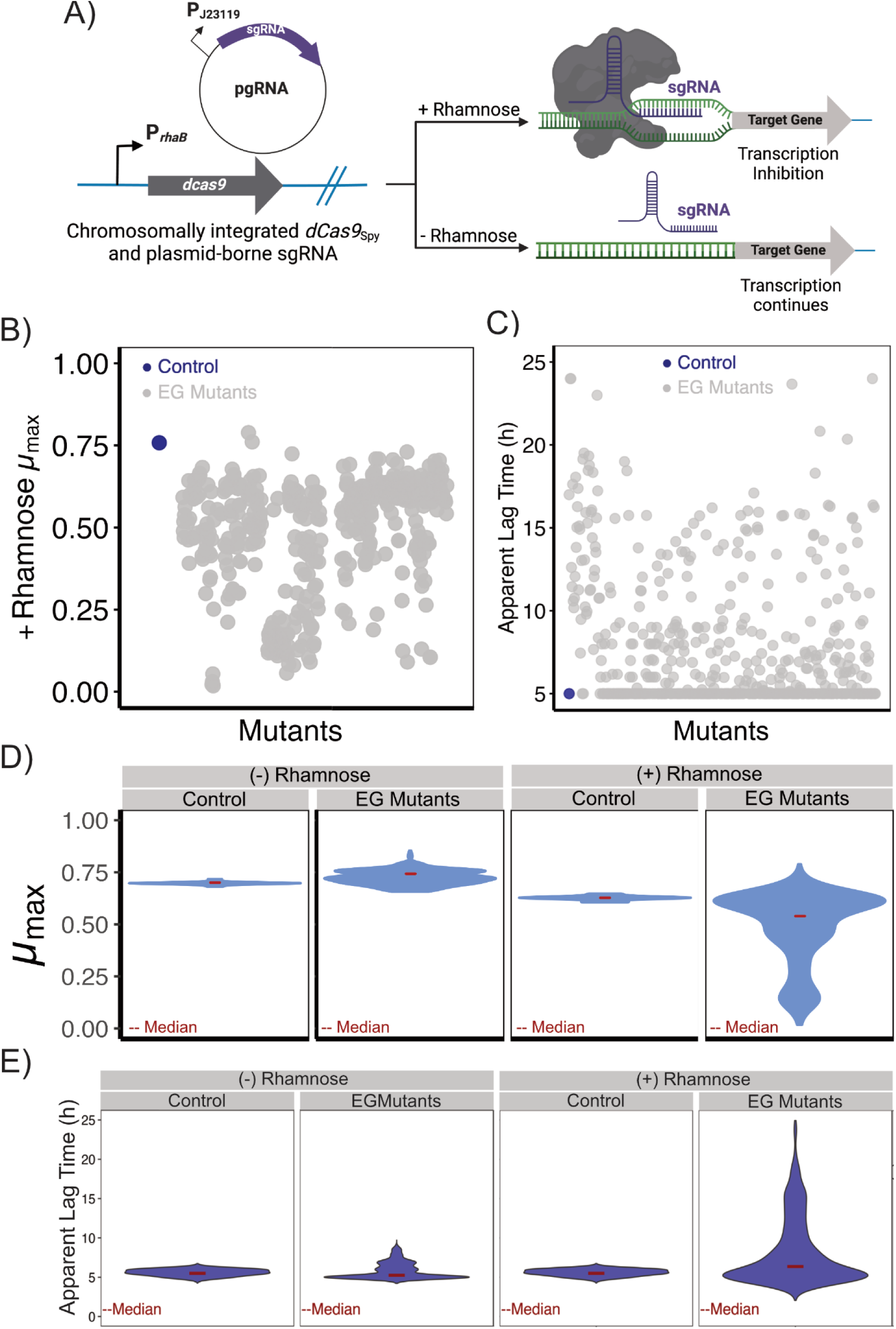
Construction and characterization of an arrayed essential gene mutant library (EGML) using CRISPRi. A) The EGML was constructed based on a dCas9_Spy_ based CRISPRi system previously developed for *Burkholderia*. The system consists of a chromosomally integrated *dCas9* under a rhamnose-inducible promoter and plasmid-borne sgRNA. The image was reconstructed from Hogan et al.^32^ and created with Biorender. B-E) Maximum growth rate (µ_max_) (B and D) and apparent lag time (C and E) of the individual essential gene knockdown mutants and their distribution in the presence of rhamnose. The blue dot indicates the pgRNA-nontarget control, and the grey dots indicate individual essential gene mutants.

Gene silencing with CRISPRi is more efficient when transcription initiation is blocked by targeting the promoter region in the non-template strand^37,38^. As the transcription start sites for most *B. cenocepacia* essential operons are unknown, we evaluated whether sgRNAs could be designed based on the proximity to the translation start sites. Fifteen essential genes from different Cluster of Orthologous Genes (COGs) categories were targeted with sgRNAs designed at different distances from their translation start sites (**Supplementary Fig. 2A**; **Supplementary Table 3**). These sgRNAs were cloned into the sgRNA expressing vector, pSCB2-gRNA, and introduced into *B. cenocepacia* K56-2::*dCas9* via tri-parental mating ^32^. We observed defective growth phenotypes for all targeted genes (**Supplementary Fig. 2B**), in agreement with recent findings^6,14,32^. To design sgRNAs targeting all essential genes, we developed a custom sgRNA design pipeline **(Supplementary Fig. 3).** While computational tools for sgRNA design exist^39–46^, they primarily cater to eukaryotic genomes and have not been used in GC-rich organisms like *Burkholderia*. By combining the insights from our pilot study and published protocols^6,7,17,37,38^, we designed 615 sgRNAs targeting the *B. cenocepacia* K56-2 essential genome^33,34^ and one sgRNA as a non-targeting control **(Supplementary Fig. 3; Supplementary Table 4**).

To test the efficiency of sgRNAs and evaluate the growth phenotype of CRISPRi silencing of essential genes, mutants were grown clonally in microtiter plate format. From this, maximum growth rate (µ_max_) (**Supplementary Fig. 4A)** and apparent lag time (**Fig. 1B, C**) were calculated. In non-inducing conditions, mutants and control exhibited similar growth parameters (median µ_max_ of 0.7h^-1^ and 0.73h^-1^, and apparent lag time of 5.1h and 5.2h, respectively (**Fig. 1D, E**-rhamnose). Upon induction of the CRISPRi system, mutants showed a deviation towards a lower µ_max_ and a more extensive apparent lag time (**Fig. 1D, E** +rhamnose). We arbitrarily designated mutants with a growth defective phenotype when they observed a 15% reduction in µ_max_ or a 15% longer apparent lag time in inducing conditions compared to the pgRNA-nontarget control. We obtained mutants with an observable growth defect in 92% of the essential gene targets.

To determine whether general essential functions can explain the observed variation in growth phenotypes, we sorted the mutants based on the COG categories and μ_max_in the presence of rhamnose. We observed the slowest µ_max_ of the CRISPRi knockdown mutants targeting translation processes (COG J and O), membrane biogenesis (COG M) and DNA replication and repair (COG L) compared to the pgRNA-nontarget control (p <1×10^-5^, t-test; **Supplementary Fig. 4B**). 80% of the knockdown mutants had at least 15% longer apparent lag times compared to the pgRNA-nontarget control (**Fig. 1C, E**). Protein synthesis and cell wall-related mutants had the longest apparent lag time (**Supplementary Fig. 4B**), and some of the mutants never reached the exponential phase (**Supplementary Fig. 4A**). Together, the EGML growth phenotypes were very diverse, highlighting the differing essential functions under CRISPRi control. This diversity poses a challenge to chemical-genetic profiling using a pooled library as the extent of target gene knockdown may vary across the mutants ^47,48^.

### Mutant depletion during pooled growth is explained by the combination of clonal µ_max_ and lag time

Three important considerations for chemical-genetic profiling using pooled essential gene libraries are that i) individual knockdown depletion during pooled growth must be accurately detected, ii) the library must represent the essential genome, and iii) sensitizing conditions must be achieved, where target downregulation effectively reports the mechanism of action of the compound under investigation, Using sgRNA coding sequences as unique barcodes for Illumina sequencing, we obtained accurate detection of individual mutant depletion from artificially depleted mutant pools. We observed a strong correlation between biological replicates of CRISPRi-Seq with small (50-member) and large (150-member) mutant pools, confirming the method’s ability to detect changes in relative abundance accurately (**Supplementary Fig. 5**) and demonstrating its reliability (**Supplementary Fig. 6**). The next step was to obtain a pooled EGML representing the essential genome in sensitized conditions. In our experience, knockdown mutants partially depleted in essential gene products are specifically sensitized to their cognate growth inhibitors when their clonal growth is reduced between 30 and 60% with respect to the wild type^47,48^. A stronger growth reduction tends to cause general sensitivity to many stressful conditions. Because competitive growth enhances knockdown mutant sensitivity^47^, we reasoned sensitization during pooled growth could be achieved at less than 50% depletion levels. To initially characterize pooled growth, the EGML strains were equally combined in a pooled inoculum (1X pool), allowed to grow at different rhamnose concentrations (**Supplementary Fig. 7)** and their relative abundance was determined by CRISPRi-Seq. Non-inducing conditions (likely causing basal levels of dCas9 expression) were sufficient for reducing fitness (depletion above 25%) in 77% of the pooled CRISPRi mutants (**Supplementary Fig. 8A**). However, even in the mildest conditions, the pooled EGML contained mutants representing 120 essential targets that were depleted by more than 50%.

To obtain a pooled EGML with representation of the essential genome in sensitized conditions, we reasoned that we could balance each mutant’s growth and sensitization levels by altering the inoculum. While the growth rate of the knockdown mutants (µ_max_) inversely correlated with the lag time (**Fig. 2A**), neither the apparent lag time nor µ_max_ of individual mutants in clonal growth could serve as predictors of depletion levels during pooled growth (**Fig. 2B, C**). Similarly, the end-point maximum growth inhibition did not correlate with depletion during pooled growth (**Fig. 2D**). We then reasoned that a combination of the individual growth parameters and mutant fitness due to competition would impact mutant depletion levels during pooled growth. To test this possibility, we used multivariate polynomial regression to model the relationship between the clonal features (µ_max(+Rha)_ and apparent lag time) and depletion levels during pooled growth (**Supplementary Note 2)**. Multivariate polynomial regression extends polynomial regression to multiple independent variables, capturing complex non-linear relationships. The fitted model predicted depletion scores based on the relationship between the clonal features. We observed a strong correlation between the experimentally observed and predicted mutant depletion (**Fig. 2E**). Our analysis demonstrates that depletion levels during pooled growth can be predicted by combining CRISPRi mutants’ clonal growth parameters.

**Fig. 2:**
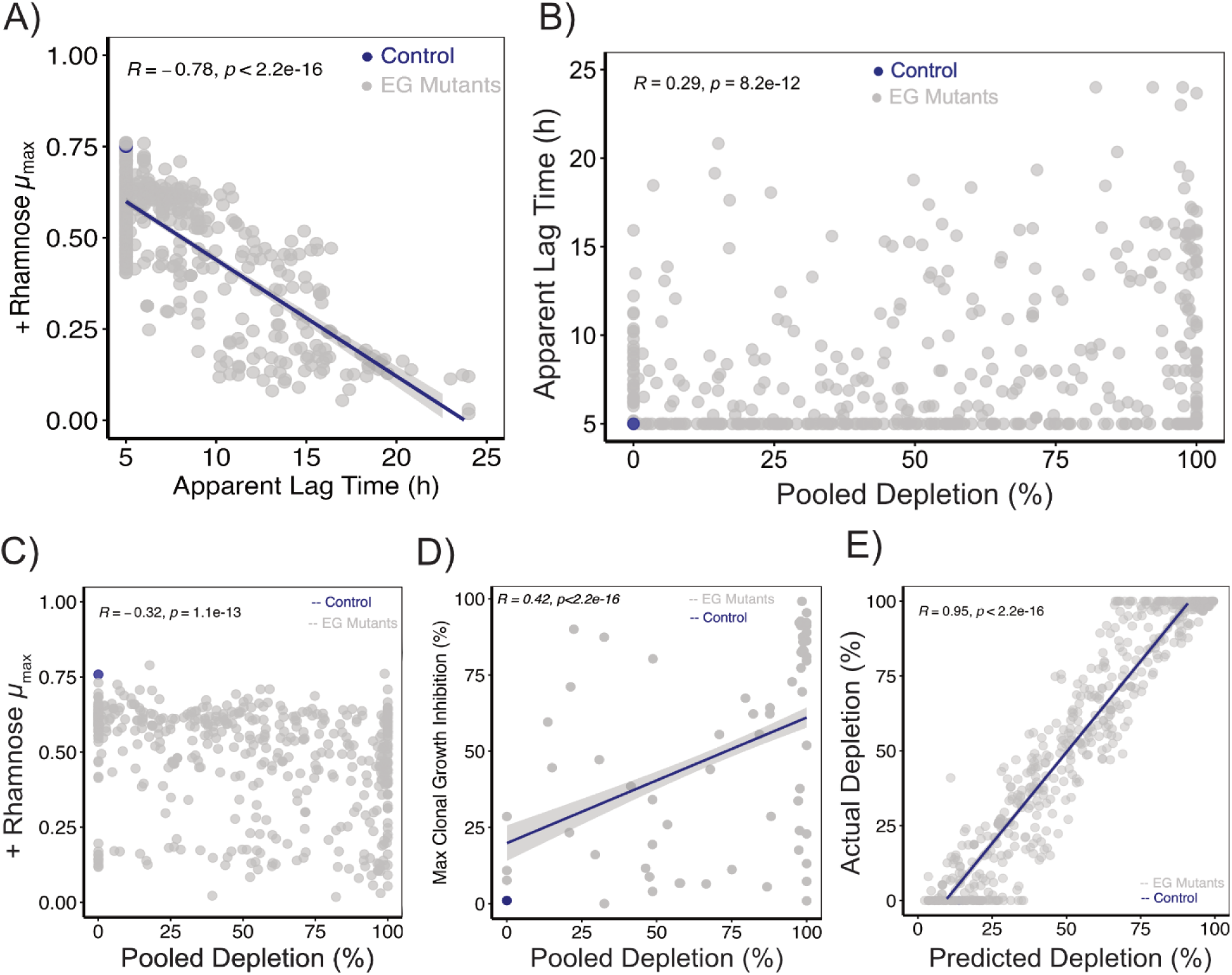
EGML strain depletion during pooled growth is related to the clonal growth rate (µ_max_) and apparent lag time. A) Correlation of clonal μ_max_ with apparent lag time of the mutant strains in the presence of rhamnose. B-C) Correlation of clonal apparent lag time (B) and μ_max_ (C) with the relative percent depletion of the mutants from the 1X (equal) pool. D) Comparison of the maximum percent clonal growth inhibition against the depletion from 1X pool during pooled growth of the mutants. Data shown for 40 mutants. E) Comparison of predicted and experimentally derived mutant depletion from the 1X pool. Predicted mutant depletion was calculated based on the clonal growth characteristics (µ_max (+Rha)_ and apparent lag time). The correlation coefficient was calculated based on the Pearson method. The mutant relative abundance in the pools was calculated based on the proportions of reads for the mutants in a pool and then compared to a control pool that had not been grown after pooling. Results are the mean of two independent biological replicates. The blue dot indicates the non-targeting control, and each grey dot indicates an individual mutant strain. Blue lines in panels A, D and E indicate the regression lines.

### Rational design and construction of CIMPLE

The predicted depletion of CRISPRi mutants during pooled growth indicated that a pooled EGML would lack a complete representation of the *B. cenocepacia* K56-2 essential genome due to particular mutants falling below the detection threshold (**Fig 2E**). We reasoned that manipulating the relative abundance of the CRISPRi mutants’ inocula could change the competition dynamics, increasing the number of sensitized mutants to achieve coverage of the essential genome in the final pool (**Fig. 3A**). In this scenario, mutants predicted to be depleted at different degrees (’sick and highly sick mutants’) could be inoculated at different proportions (spiked pool) into distinct pools. Therefore, the initial inoculum of the mutants was arbitrarily scaled based on the mutants’ expected depletion from the equal EGML (1X pool). Mutants expected to be depleted more than 75% (highly sick) in the equal EGML were inoculated at ratios of 5X, 10X, or 20X. Mutants with 60-75% depletion (‘sick’) were inoculated at ratios of 3X, 5X, or 10X, while mutants with 50-60% depletion (‘sick’) were inoculated at ratios of 2X, 3X, or 5X, (**Supplementary Table 5**) resulting in three differentially-inoculated pools: 5X3X2X, 10X5X3X and 20X10X5X pools. Mutants with expected depletion below 50% were pooled at 1X. We observed undesired mutant overrepresentation after growth from the 10X5X3X and 20X10X5X pools (**Supplementary Fig. 8B-C).** On the other hand, the 5X3X2X-inoculated pool demonstrated the highest proportion of sensitized mutants as most (∼89%) of the mutants were depleted less than 50% (**Fig. 3B**). While 67 mutants became highly abundant (enriched) in the 5X3X2X pool (compared to 57 in the equal EGML), pooled growth from the 5X3X2X-inoculated pool provided ∼90% coverage of the *B. cenocepacia* K56-2 essential genome in one single sensitizing condition (non-inducing) (**Supplementary Fig. 8D**). In contrast to the previous approaches with handling individual mutants^7^ or multiple pools^3^, our approach allowed representation of nearly the entire essential genome in one single condition, which can streamline and increase the throughput of chemical-genetic profiling experiments.

**Fig. 3:**
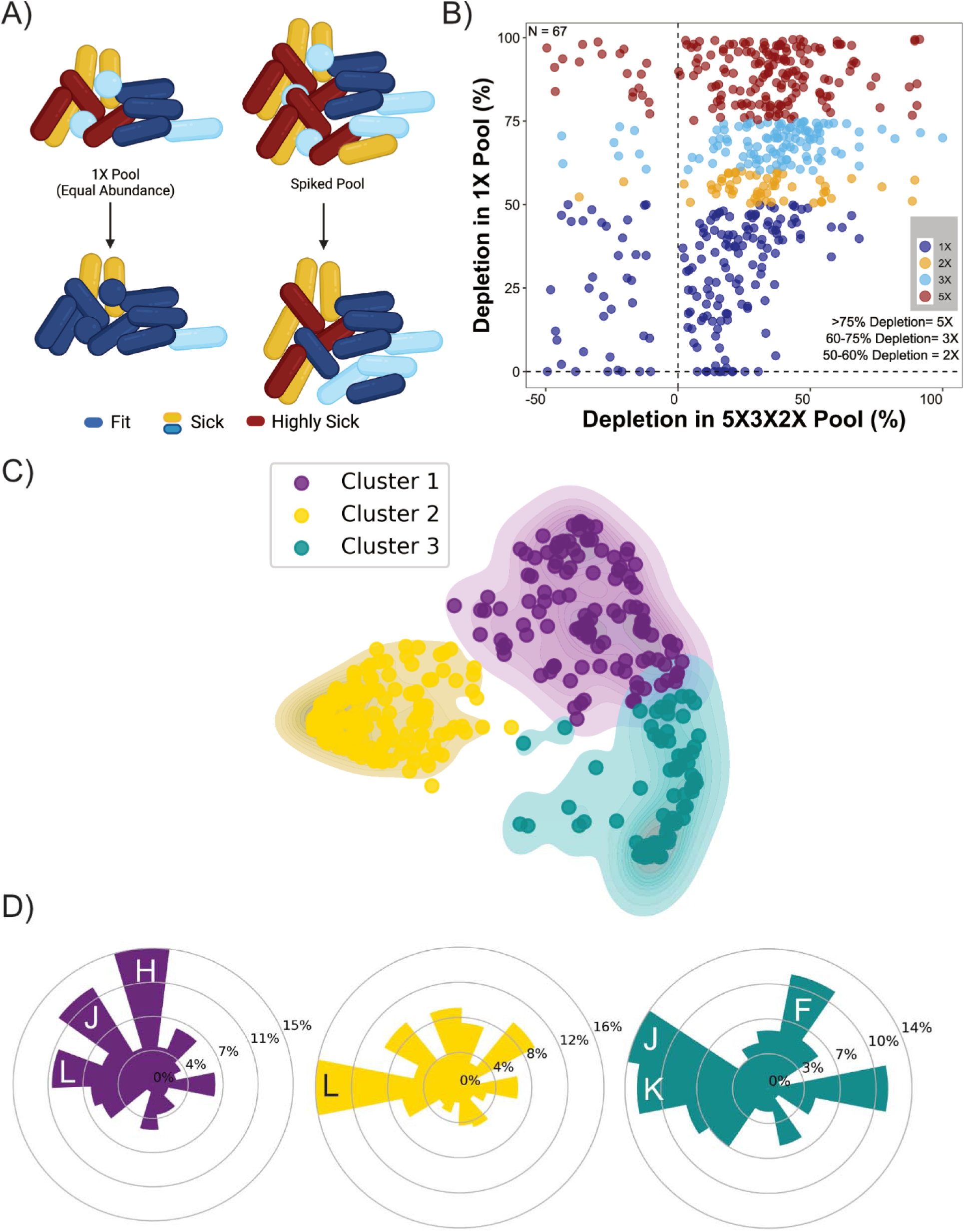
Spiking of the sick EGML strains enables recovery of the mutants in the spiked pool. A) Schematic illustrating the relative fitness of the CRISPRi mutants in the 1X and spiked pool. Increasing the initial relative abundance of the highly sick mutants conferred a competitive advantage, enhancing their fitness. B) Comparison of mutant depletion between the 1X (equal) and 5X3X2X spiked pools. The mutant relative abundance (RA) in the pools was calculated based on the proportions of reads for the mutants in a pool and then compared to a control pool that had not been grown after pooling. Negative depletion indicates enrichment of the mutants compared to a control pool that had not been grown after pooling. Results are the mean of two independent biological replicates. C) Kernel PCA with K-means clustering of the mutants based on the clonal characteristics (µ_max_ and apparent lag time) and percent depletion from the pool. D) Percent of COGs in each cluster from panel C.

To investigate the possible link between fitness during pooled growth and essential functions, we weighed the contribution of different parameters by a principal component analysis (PCA) and compared the obtained cluster of mutants with their COG categories. To that end, we used a Kernel PCA^49^ combined with K-means clustering^50^ and clustered the mutants based on µ_max_, apparent lag time, mutants’ pooled fitness and fold inoculum level in the 5X3X2X-inoculated pool. We clustered the mutants into three clusters based on K-means clustering and projected the mutants in two-dimensional PCA space (**Fig. 3C)**. Mutants that were inoculated at 1X proportions comprised 98% of the mutants grouped in cluster 2, which was specifically enriched in the COG category DNA replication and repair (L) (**Fig. 3D**). On the other hand, mutants that were depleted more than 75% constituted 91% of cluster 3, which was enriched in translation (J), nucleotide metabolism and transport (F) and transcription (K). In contrast, cluster 1 was more diverse as only 50% of the mutants belonged to the group of mutants inoculated 2X and 3X (**Fig. 3D**). This cluster was also functionally diverse, containing the COG categories coenzyme metabolism (H), translation (J) and DNA replication and repair (L).

Overall, functional diversity of essential functions partially explained differential fitness during competitive growth. However, the prediction of depletion levels from clonal mutant growth guided a rationally designed pooled inoculum that changed the competition dynamics during pooled growth, increasing genomic coverage of essential genes and expected levels of sensitization. This approach enabled the creation of an optimized <u>C</u>RISPR interference-<u>m</u>ediated <u>p</u>ooled library of <u>e</u>ssential genes (CIMPLE) in *B. cenocepacia* K56-2.

### CIMPLE with CRISPRi-Seq (CIMPLE-Seq) captures drug-target interactions and the targets of known antimicrobials

Conventional pooled libraries, although powerful, can fail due to reduced effective genomic coverage when unequal fitness of mutant strains before drug exposure causes mutants to drop below the detection threshold in the pool. In contrast, CIMPLE-Seq allows essential genome coverage to report antibiotic mechanisms of action. For instance, the *dcw* operon (containing *ftsZ*) and *gyrB* were not detected in the 1X pool (100% depleted); however, by changing the corresponding mutants’ relative abundance in the 5X3X2X inoculum, the *dcw* operon and *gyrB* genes were represented in CIMPLE (**Supplementary Fig. 9**). To confirm that CIMPLE-Seq can detect the depletion of specific mutants when treated with their cognate antibiotics, we exposed CIMPLE to a subinhibitory concentration (IC_30_) of antibiotics from different classes with known cellular targets (**Fig. 4A**; **Supplementary Table 6**). We measured the relative abundance of the CRISPRi mutants from the pool in each treated condition and compared that to the DMSO control. The normalized relative abundance was highly correlated between the independent biological replicates (**Supplementary Fig. 10**). We focused on the significant chemical-genetic interactions between compounds and essential gene mutants (**Fig. 4B-E**). When we exposed CIMPLE to C109, a benzothiadiazole compound that targets FtsZ^48^, we observed a significant depletion of the CRISPRi mutant targeting the division and cell wall cluster (**Fig. 4B**; *dcw* cluster; K562_RS16765; to K562_RS16835), which contains *ftsZ*. The *dcw* is an essential operon that maintains cell division and cell wall synthesis in a regulated manner, starting with FtsA and FtsZ^51^. While we observed seven significantly depleted hits in the absence of rhamnose (including the specific target *dcw* cluster), the number of hits was higher in the presence of rhamnose (0.1%) which still included the CRISPRi mutants targeting the *dcw* operon (**Fig. 4C**). Likewise, both *gyrA* and *gyrB* targeting CRISPRi mutants were significantly depleted in the presence of novobiocin (**Fig. 4D-E**). Novobiocin is an aminocoumarin antibiotic that binds to the ATP binding site of GyrB (which acts by forming a complex with GyrA) and inhibits enzymatic activity by impeding ATP hydrolysis^52^. Novobiocin also caused significant depletion of DNA polymerase III subunit β (*dnaN*; K562_RS02185), transcription termination factor Rho (*rho*; K562_RS09165), Holliday junction resolvase (*ruvC*; K562_RS16145), a hypothetical protein (K562_RS11950), and UDP-3-O-[3-hydroxymyristoyl] N-acetylglucosamine deacetylase (*lpxC*; K562_RS16755). LpxC catalyzes the deacetylation of UDP-3-O-acyl-N-acetylglucosamine, which is a crucial step in the synthesis of lipid A. As a lipopolysaccharide (LPS) component, lipid A plays a critical role in the structural integrity of the outer membrane. Overall, we found that non-inducing conditions were sufficient to capture an antibiotic susceptibility profile and to identify the specific cellular target.

**Fig. 4:**
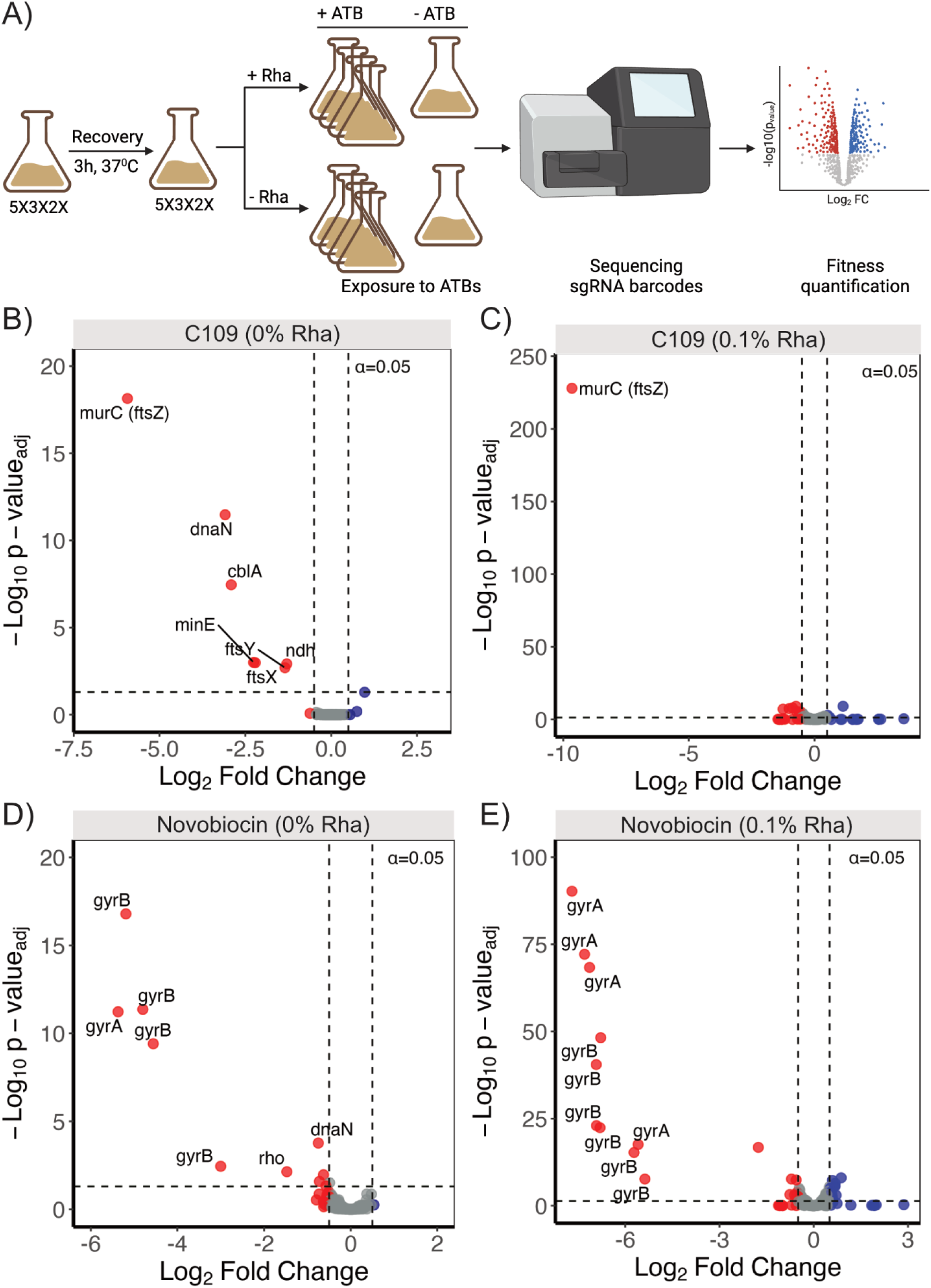
CRISPRi-Seq identifies known and novel drug-target interactions. A) Overview of the CRISPRi-Seq workflow. Exposure of the CIMPLE pool to subinhibitory concentration (IC_30_) of C109 (B-C) and novobiocin (D-E). CIMPLE pool exposure to antibiotics without (B and D) and with (C and E) of 0.1% rhamnose. Blue and red dots represent enriched and depleted mutants (B-E), respectively. The horizontal dashed line indicates the significant threshold (α = 0.05) (B-E). Vertical dashed lines indicate a log_2_ fold change of ±0.5. Results are the mean of two independent biological replicates.

### CIMPLE-Seq identified peptidyl-tRNA hydrolase (Pth) as a target of PHAR(A)

We previously employed a deep learning model to predict bacterial growth inhibition in a natural product library containing over 224,000 compounds^53^. Our approach led to the identification of STL529920 (renamed to PHAR(A) herein), a growth inhibitory compound with a chemical scaffold not previously linked to antimicrobial activity^53^. PHAR(A) exhibited broad growth inhibitory, albeit weak, activity against *B. cenocepacia* K56-2 and the ESKAPE pathogens^53^ by an unknown mechanism. To elucidate PHAR(A)’s mechanism of action, we exposed CIMPLE to 50µM of PHAR(A) either in non-inducing conditions or with 0.1% rhamnose. In the absence of rhamnose, CIMPLE-Seq revealed significant depletion of a CRISPRi mutant targeting the peptidyl-tRNA hydrolase (Pth; K562_RS04080), as well as knockdown mutants of translation-related genes 30S ribosomal protein S1 (*rpsA*; K562_RS14090) and 50S ribosomal protein L20 (*rplT*; K562_RS11325) (**Fig. 5A**). In the presence of 0.1% rhamnose, additional knockdown mutants of genes related to translation (a hypothetical protein (K562_RS03150), aminopeptidase P (*pepP*; K562_RS16120), tRNA-specific 2-thiouridylase MnmA (*mnmA*; K562_RS16105)) and cell envelope related protein (WaaA (*waaA*; K562_RS14990)) were significantly depleted from the pool (**Fig. 5B)**.

**Fig. 5:**
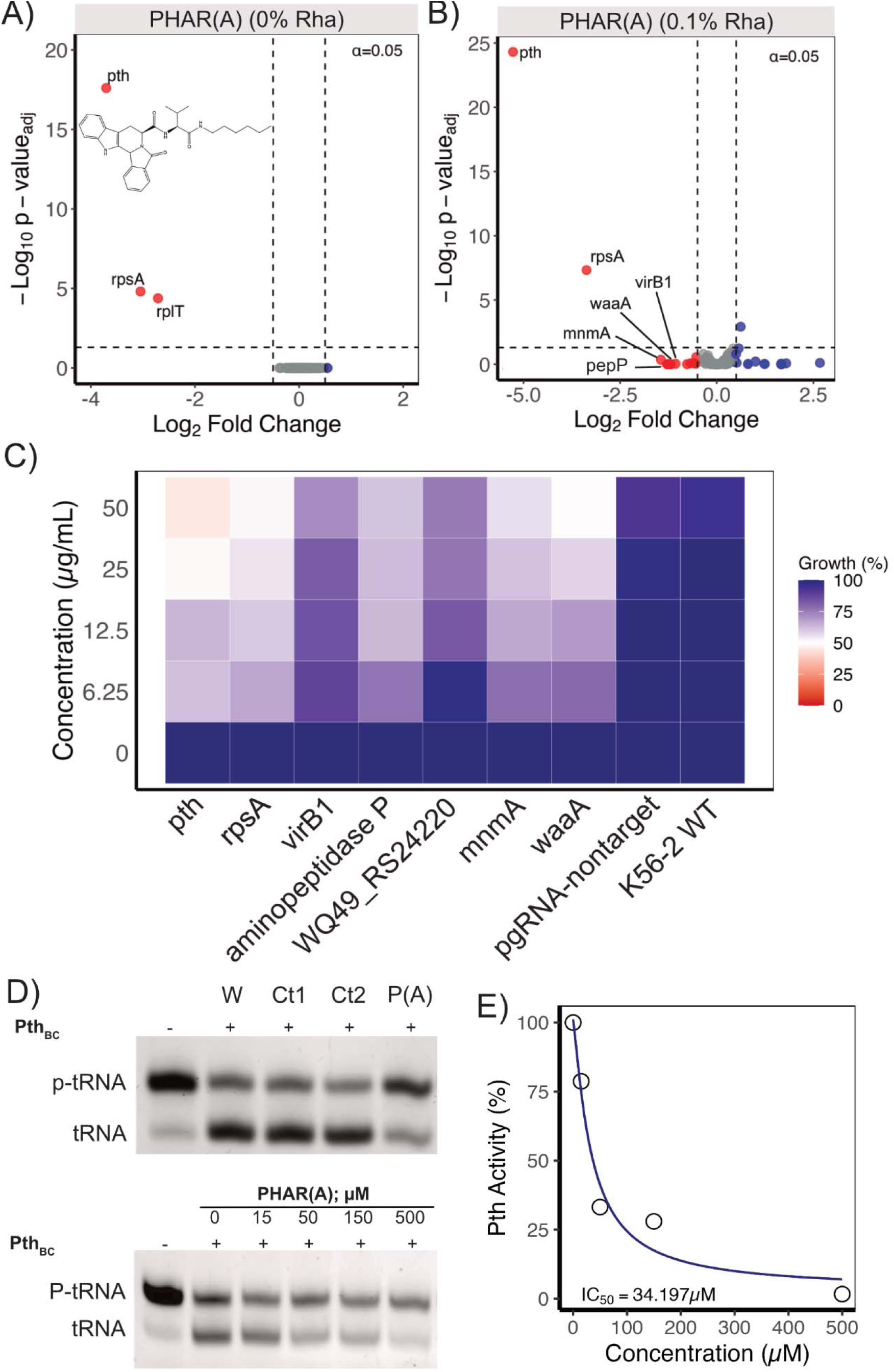
PHAR(A) inhibits bacterial growth by impeding Pth activity. A-B) Exposure of the CIMPLE library to 50µM PHAR(A) without (A) and with (B) 0.1% rhamnose. C) Exposure of a Pth CRISPRi mutant exhibited hypersensitivity to PHAR(A). D) *In vitro* inhibition of *B. cenocepacia* K56-2 Pth (Pth_BC_) activity as shown by migration of intact peptidyl-tRNA (p-tRNA) and cleaved (t-RNA). Samples were electrophoretically separated in 12% polyacrylamide gel with 7M urea and stained with ethidium bromide. Water, Control compounds, and PHAR(A) are abbreviated as W, Ct and P(A), respectively. E) Quantification of percent Pth activity based on the band intensity (in panel D) of the cleaved tRNA relative to the (-)Pth control. The band intensity was measured using ImageJ^77^, and the IC_50_ was calculated using the “DRC”^78^ package in R.

The abundance of ribosome-associated hits suggested that PHAR(A) may interfere with the translation machinery. As Pth was the strongest hit, we investigated if PHAR(A) inhibits Pth, a highly conserved bacterial essential protein required for recycling stalled ribosomes^54^. To confirm the chemical-genetic interactions detected by CRISPRi-Seq, we re-assessed PHAR(A) susceptibility of clonally grown CRISPRi mutants in sensitizing conditions (rhamnose concentration to suppress growth by 30%, RhaIC_30_) (**Supplementary Table 7**). The Pth knockdown mutant was the most susceptible to PHAR(A) compared to the non-targeting CRISPRi control (**Fig. 5C)**, resembling the CIMPLE-Seq profiling of PHAR(A). To lower levels, the other translation (RpsA, MnmA, PepP) and secretion system-related knockdown mutants (VirB1) were also sensitive in the presence of PHAR(A). To investigate if Pth is inhibited by PHAR(A), we purified *B. cenocepacia* K56-2 Pth (Pth_BC_). We incubated the purified Pth_BC_ with peptidyl-tRNA to assess the Pth-mediated peptidyl-tRNA cleavage activity in the presence and absence of PHAR(A). We observed a dose-dependent inhibition of the recombinant Pth_BC_ activity by PHAR(A) starting from 15µM (PHAR(A) IC_50_ = 34.2µM; (**Fig. 5D-E**)), but not by PHAR(A) analogs that did not have any growth inhibitory activity (**Supplementary Fig. 11**)^53^. To identify the potential PHAR(A) binding site, we performed a molecular docking analysis. Docking of PHAR(A) into Pth_BC_ showed that PHAR(A) might interact with residues that were reported previously to be critical for the catalysis (Asn10, Asp93, Asn114 and the essential His20)^55,56^ and substrate binding (Asn10, His20, Asp93, and His113, Asn114)^55–59^ (**Fig. 6A**). While the inactive analogs of PHAR(A) (Ct1 and Ct2) are predicted to interact with these residues (**Fig. 6B-C**), they appear to lack interactions with Asn10, indicating the critical function of Asn10 for catalytic activity. Indeed, site-directed mutagenesis of Asn10 residue leads to a 200-fold decrease in Pth activity^56^. These results indicate that PHAR(A) is a specific inhibitor of the bacterial Pth enzymatic activity and highlight the utility of CIMPLE-Seq to identify both direct and indirect drug-target interactions involved in the mechanism of action of novel bioactive compounds.

**Fig. 6:**
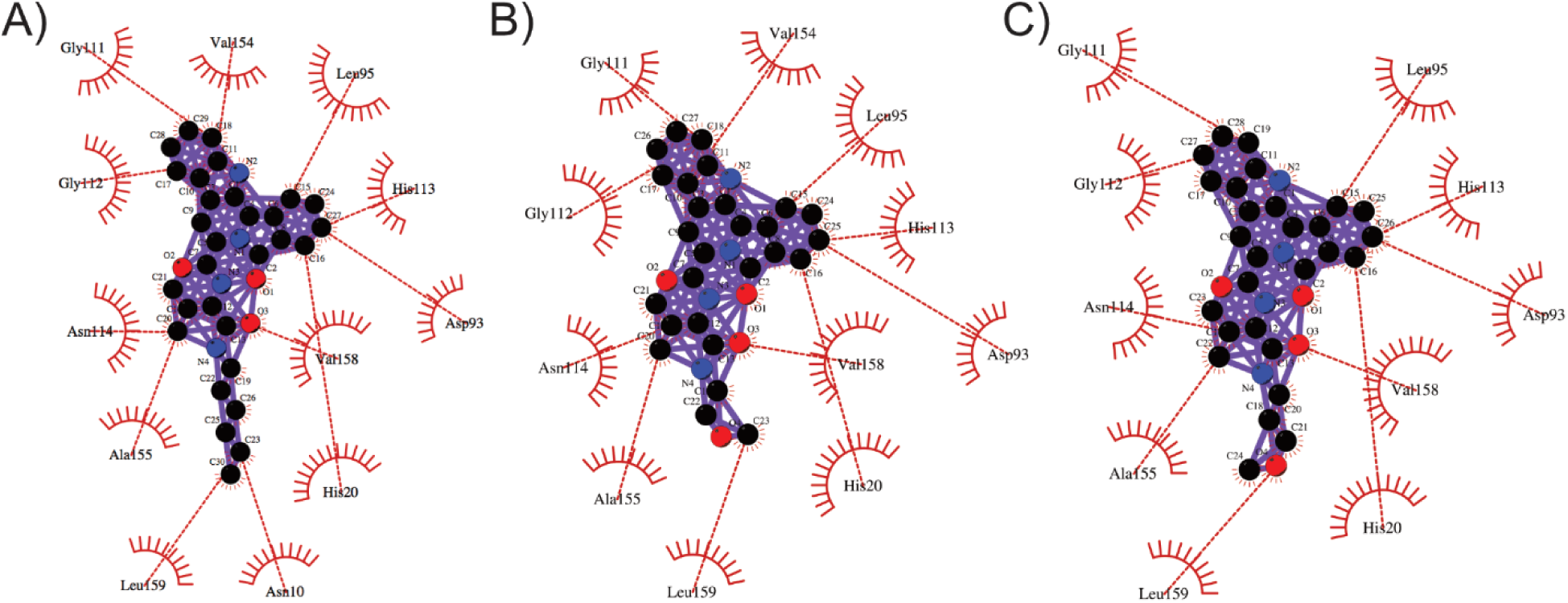
PHAR(A) may interact with amino acid residues critical for Pth enzymatic and binding activity. A-C) Interactions of PHAR(A) (A), control compound 1 (B) and control compound 1 (C) with *B. cenocepacia* K56-2 Pth (Pth_BC_). The spoked arcs represent protein residues, and the dashed red line represents hydrophobic interactions. The most favourable docked model is shown. The red, blue and black circles within the ligands (compounds) represent oxygen, nitrogen, and carbon atoms, respectively. Pth_BC_ 3D structure was generated using Alphafold. Docking was performed with Autodock vina within Python prescription (PyRx). The plots were generated using LigPlot^+^.

## Discussion

Essential gene mutant libraries are valuable tools for genetic interaction mapping to identify the mechanism of action of novel antibiotics. Here, we have constructed a comprehensive, arrayed essential gene knockdown mutant library targeting each of the identified *B. cenocepacia* K56-2 essential genes. Conventional pooled libraries risk incomplete coverage of the targeted essential genome, potentially compromising drug target identification and downstream analysis. Considering the mutants’ growth when grown clonally and in a competitive pooled growth environment, we created an optimized CRISPRi-mediated pooled library of essential genes (CIMPLE) by predicting pooled depletion levels from clonal growth parameters and changing the relative abundance of the mutants in the pooled inoculum. We demonstrated the utility of CIMPLE-Seq in identifying the mechanism of action of known and novel bioactive compounds.

CRISPRi-based libraries offer distinct advantages over libraries created with other genetic tools. Here, we exploited the ability of CRISPRi to titrate essential gene expression. However, a portion of the mutants did not show the expected growth defect in clonal growth environments upon dCas9 induction (**Supplementary Fig. 4A**; **Fig. 2A-B**). Possible reasons are insufficient target gene silencing by the CRIPSRi machinery, high target protein turnover rate, and low required levels of the target protein. However, we observed depletion of those mutants during pooled growth (**Fig. 2A-B**, **Supplementary Fig. 9A**), indicating that the competitive pooled growth exacerbated subtle mutant phenotypes. Although no clear linear correlation was observed between the pooled depletion and the apparent lag time or clonal growth rate when examined individually (**Fig. 2A-B**), our multivariate polynomial approach enabled us to discern the interplay between the mutants’ depletion during pooled growth and a combination of their clonal growth characteristics. (**Fig. 2E**).

We first benchmarked CIMPLE-Seq by identifying the known drug-target interactions of GyrAB and FtsZ with novobiocin and C109, respectively. CIMPLE was instrumental in representing the highly sick mutants (in the pool) that would have otherwise been missed in the 1X library. For example, the *dcw* operon (containing *ftsZ*), *gyrB*, and *dnaN* mutants were outcompeted to extinction in the 1X pool (100% depleted) (**Supplementary Fig. 9A**). FtsZ (part of the *dcw* cluster) is targeted by C109, GyrB is targeted by novobiocin, and DnaN also plays a role in novobiocin-mediated sensitivity. By artificially altering their initial relative abundance in the CIMPLE library, we ensured their inclusion, allowing us to capture and reaffirm the mechanisms of action of C109 and novobiocin (**Supplementary Fig. 9B**). Our study also identified the involvement of DnaN, Rho, LpxC and K562_RS11950 as potential novel mediators of novobiocin sensitivity. These findings suggest a complex mechanism for novobiocin’s efficacy, involving potential interactions with essential cellular components (DnaN and Rho) and membrane-associated proteins (LpxC and K562_RS11950).

Next, we used our approach to study the antimicrobial mechanism of PHAR(A), which was attractive due to its unique chemical scaffold. We observed depletion (**Fig. 5A-B**) and clonal enhanced sensitivity (**Fig. 5D**) of knockdown mutants targeting *pth*, *rpsA*, *mnmA*, *pepP*, and *waaA*. RpsA, MnmA, Pth, and PepP are involved in protein synthesis and regulation (**Supplementary Fig. 12**). PHAR(A) inhibited the activity of Pth *in vitro*, suggesting that PHAR(A) could target Pth *in vivo*. While MnmA, RpsA, and PepP may not be the direct targets of PHAR(A), their roles in translation, tRNA modification, and peptide metabolism are interconnected^60–63^. Ribosomal protein S1 (RpsA) serves as a gatekeeper at the exit of the mRNA channel on the small ribosomal subunit, facilitating mRNA binding and initiation of translation^60^. Concurrently, MnmA modifies tRNAs, ensuring their fidelity in decoding mRNA codons, thus directly influencing the precision of protein synthesis^61^. Pth is an essential bacterial enzyme that cleaves the ester bond between the tRNA and the peptide after peptidyl-tRNA is accidentally dropped off from the ribosome during protein synthesis, essentially recycling the tRNAs for reuse in translation^63^. Complementing this process, PepP degrades surplus peptides into amino acids, contributing to cellular recycling and maintaining metabolic equilibrium^62^. The reasons for Pth essentiality are still unclear, but this enzyme, besides a well-established function in tRNA recycling, has been recently shown to play a pivotal role in stress response and ribosome quality control^64^. Therefore, inhibition of Pth may result in the accumulation of peptidyl-tRNAs associated with stalled ribosomes in the cell, which, in turn, might inhibit protein synthesis by decreasing not only the overall amount of deacylated tRNAs but also free 50S subunits available for translation. Thus, dysregulation of translation induced by Pth inhibition can lead to cellular stress, altered metabolism, and cell death.

While the reasons why *waaA* downregulation enhanced the activity of PHAR(A) were not further investigated, increased permeation through the cell envelope was likely involved. *waaA* encodes a transferase involved in the biosynthesis of lipopolysaccharides (LPS) in gram-negative bacteria. WaaA catalyzes the addition of an acyl chain to the lipid A portion of (LPS)^65^. This acylation helps stabilize the outer membrane and contributes to its impermeability^66^. The absence of functional WaaA and the consequent truncation of LPS can enhance the permeability of the bacterial outer membrane, enhancing their susceptibility to antibiotics^65,67^.

Although PHAR(A) is a specific inhibitor of Pth, it exhibited relatively weak inhibitory activity. Our molecular docking analysis suggests that the aliphatic tail of PHAR(A) might be critical for interactions with key residue (Asn10) involved in catalytic activity (**Fig. 6**). This observation is consistent with our previous screening of 15 unique PHAR(A) analogs, showing that lack of the hydrocarbon tail correlates with inactivity of a compound^53^. Structural characterization of Pth in complex with PHAR(A) combined with a structure-activity relationship study could provide valuable insights into the molecular interactions for rational design of PHAR(A) analogs with stronger antibacterial activity.

While CIMPLE-Seq provides a valuable tool for identifying potential drug-target interactions, we acknowledge the following limitations. Firstly, the 67 mutants present in CIMPLE at relatively high abundance may limit the detection of weak or indirect drug-target interactions specific to those targets. We expect this to be minimized by reiterating the relative abundance of mutants in the CIMPLE library. Secondly, the polar effect of CRISPRi, which represses transcription of downstream genes within the same operon, may complicate the interpretation of drug-target interactions, potentially obscuring the direct target(s)^6^. The putative drug(s) can be validated with follow-up experiments with our arrayed library, partially mitigating the problem.

## Methods

### Data Reporting

No statistical methods were used to predetermine sample size. The experiments were not randomized, and investigators were not blinded to allocation during experiments and outcome assessment.

### Bacterial strains and growth conditions

Bacterial strains used in this study are provided in **Supplementary Table 8**. All strains were grown in LB-Lennox medium (Difco) at 37°C with continuous shaking at 230rpm unless otherwise specified. The following selective antibiotics were used: trimethoprim (Sigma; 50 µg/mL for *E. coli* strains and 100 µg/mL for *B. cenocepacia* strains). All plasmids used in this study are listed in **Supplementary Table 9**. L-rhamnose (Sigma) stock solution was prepared to 20% in de-ionized water and then filter sterilized.

### Designing the sgRNAs

The sgRNAs were designed using a custom Python script that produces an output file containing ready-to-order primers to clone the sgRNAs. Briefly, the script splits the genome (in Genbank format) into two vectors (forward and reverse strand) and extract the sequence based on a provided query range. For example, if the script has a query range of −100 to +50 for a given gene, the script will extract 150bp, 100bp upstream and 50bp downstream of the translation start site (ATG). The script will then look for the possible sgRNAs based on the availability of the PAM sites and extract the candidate sgRNAs. The candidate 20bp sgRNAs are then mapped to the genome to determine the specificity by paring one base at a time from the 3’ end up to 12bp and will select the one sgRNA per gene that does not have off-target binding sites, is free of homopolymer stretches (≥ 5) and closest to the ATG site. The script is customizable and can be modified to target the whole genome or only the essential genome, alter the query range, and provide more than one sgRNA per gene, if necessary. The provided sgRNAs are 20bp long and are then concatenated with the sgRNA scaffold sequence to predict the secondary structure folding of the hairpin required for dCas9 binding using Quikfold algorithm from the UNAFold package (http://unafold.rna.albany.edu/?q=DINAMelt/Quickfold). Successful sgRNAs are then concatenated with the pre-designed forward primer as a 5’ extension and used for cloning the sgRNAs (see below). The operon information to select the targets was collected from the DOOR database^35^ and manually adjusted based on the transcriptomic data from Sass *et al.* ^36^. The *B. cenocepacia* K56-2 genome sequence (RefSeq accession: GCF_014357995.1) and the operon file were used as input for the Python script to design the sgRNAs. The sgRNA sequences are provided in **Supplementary Table 4**.

### Cloning the sgRNAs and construction of the CRISPRi EGML

The CRISPRi mutants were created in an arrayed manner, as previously described^32^, with modifications. Briefly, the selected sgRNAs were added as 5’-tail to a forward primer (sequence: 5’– GTTTTAGAGCTAGAAATAGCAAGTTAAAATAAGGC – 3’) that binds to the promoter region of the sgRNA expressing plasmid (pSCB2-gRNA). The resulting primers were ordered in 96-well format from IDT. All primers used in this study are listed in **Supplementary Table 10**. The sgRNA sequences were cloned into the pSCB2-gRNA by inverse PCR using reverse primer 1092 and the sgRNA-bearing primer. The PCR was performed using Q5 DNA polymerase with high-GC buffer using the following conditions: 98°C-5mins, 98°C-30s, 30 cycles (98°C-10s, 60°C-30s, 72°C-5mins), and then 72°C-2mins. The raw PCR product was used for blunt-end ligated (NEB; 30mins at 37°C). The circularized plasmids were transformed into *E. coli* DH5α chemical competent cells. PCR amplification of the sgRNAs, cloning into pSCB2-gRNA, and transformation into *E. coli* DH5α were performed in a 96-well format (**Supplementary Method**) in 96-well PCR plates (Sarstedt). The *E. coli* DH5α cells harbouring the sgRNA plasmids were delivered into a *B. cenocepacia* K56-2 mutant expressing dCas9_Spy_ (K56-2::*dCas9_Spy_*) by conjugation.

### Growth curves and clonal growth analysis of the CRISPRi EGML

The mutants were grown overnight into 96-well plates containing LB medium, selection antibiotic (trimethoprim 100µg/mL) and with/without 1% rhamnose (the dCas9_Spy_ inducer) at 37°C with continuous shaking (230 rpm). Overnight cultures were back diluted to OD_600_ of 0.01 and re-arrayed into 96-well plates in duplicate with and without 1% rhamnose. The mutants were grown at 37°C with continuous shaking at 230 rpm. Mutant growth was measured at OD_600_ in real-time using a BioTek Synergy 2 microplate reader. One hundred and five mutants showed an observable growth defect when subcultured 4 h in the presence of rhamnose. The data from the microplate reader was processed in R (version 4.3.0). Clonal maximal growth rates (µ_max_) and apparent lag times were calculated from the microplate growth curve data using ‘growth rates’ R package^68^ applying ‘easylinear’ method. This approach involves fitting linear model segments to logarithmically transformed data to calculate the µ_max_.

### Creating equal and artificially depleted pools

To create the equal and depleted pools, the mutants were individually grown overnight in 96-well format at 37°C with continuous shaking at 230 rpm. The overnight cultures were back diluted in 1:100, and the culture optical density was measured and pooled at final OD_600_ of 2.00. Glycerol was added to the pool to a final concentration of 20%. The final pool was mixed well by pipetting. All the mutants were pooled at equal abundance in the “equal pools,” and the appropriate mutants were added at lower abundances (0.5X to 100X) in the “depleted pool.” To lyse the cells for sgRNA amplification, neat DMSO was added at a final concentration of 20% (v/v) to the pools, heated at 95°C for 5 mins in a thermocycler and stored in the −80°C freezer until needed.

### Creating the CIMPLE, growth analysis and sensitizing rhamnose concentration determination

The essential gene knockdown mutants were pooled as described above to create “equal EGML”. All the mutants were pooled at equal abundances measured based on OD_600_. The “equal EGML” was grown for 4h with and without rhamnose first (to mimic the clonal sub-culturing condition) or directly grown in different rhamnose concentrations ranging from 0 to 1% for 8h (∼10–12 generations) in 96-well plates (Sarstedt) at 37°C (**Supplementary Fig. 7**). Neat DMSO was added to the pools at 20% final concentration (v/v) and treated at 95°C for 5 mins in a thermocycler and stored in the −80°C freezer. The 5X3X2X, 10X5X3X and 20X10X5X pools were created similarly, except the inoculum of selected mutants was increased from 2X to 20X as indicated. Samples were prepared for CRISPRi-Seq as described below (see *NGS library preparation and CRISPRi-Seq*).

### Multivariate polynomial regression modeling to predict the mutant depletion

Mutant depletion was estimated using a random forest regression model from ‘sklearn.ensemble’ package in Python. Depletion values were calculated based on a combination of multiple decision trees, each providing its own estimation. Clonal growth features (µ_max_ and apparent lag time in the presence of 1% rhamnose) were first transformed (feature engineered) into polynomial features using ‘PolynomialFeatures’ package in Python to capture potential non-linear relationships. A random forest regression model was fit to the transformed features and the pooled depletion data. The fitted model was applied to estimate the percent depletion values for the mutants based on the given input features.

### NGS library preparation and CRISPRi-Seq

The mutant pools (equal pools, depleted pools and the optimized EGML) were used as templates for PCR to amplify the N_20_ sgRNA region. To avoid issues related to amplification errors, the number of PCR cycles was limited to a cycle that corresponds to ∼25% maximum product. The number of cycles corresponding to ∼25% maximum product was quantified by semi-quantitative PCR (**Supplementary Fig. 14**). Illumina flow cell adaptor, Nextera index, and sequencing primer binding site were added as 5’ extension to the forward primer (sequence 5’-ACATTGACATTGTGAGCG-3’) and reverse primer (sequence primer 5’-ACTTGCTATTTCTAGCTC-3’) to enable 1-step PCR amplification. Resultant primers were used for PCR with Q5 DNA polymerase with high-GC buffer (NEB). For CRISPRi-Seq, 2 × 50µL PCR reactions were set up using the following amplification conditions: 98°C-30s, 18 cycles (98°C-10s, 52.4°C-30s, 72°C-20s), 72°C-1mins. PCR products were cleaned with SeraMag beads (Cytiva) and analyzed on a TapeStation4150 (Agilent Technologies). Amplicons were sequenced on an Illumina MiSeq for the equal and artificially depleted pools with a MiSeq reagent kit v2 micro (PE 60 bp reads) and 20% PhiX spike.

### Pilot screening of antibiotics with CRISPRi-Seq

A. *cenocepacia* K56-2 wild-type was exposed to a 2-fold gradient of different antibiotics in LB broth to determine the concentration that inhibits 30% of wild-type growth (IC_30_; **Supplemental Table. 6**). Frozen CIMPLE pool (OD_600_ 1.00; 1mL) was thawed at room temperature and added to 4mL LB broth in a 5’ glass tube. The culture was incubated at 37°C in LB medium with shaking at 230 rpm for 3 hours to recover the cells. The CIMPLE was then exposed to the IC_30_ concentration of different antibiotics (or 1% DMSO control) for 8h (∼10–12 generations) in 96-well plates (Sarstedt) in LB medium (final volume 200µL) in a humidity-controlled chamber at 37°C without shaking. 20% (final concentration v/v) DMSO was added to the wells after 8h and treated at 95°C for 5 mins in a thermocycler. The samples were prepared for CRISPRi-Seq as described above. Sequencing was performed on an Illumina MiSeq (Donnelly Centre, Toronto, Canada) with a MiSeq reagent kit v3 (PE 60bp reads), 20 dark cycles and 20% PhiX spike.

### sgRNA counts from Illumina NGS to infer mutant fitness

sgRNAs were counted from the CRISPRi-Seq data using 2FAST2Q ^69^. Since sequencing reads had a consistent structure, the locations of sgRNA sequences within each sequencing read in the demultiplexed FASTQ files (fastq.gz), along with the sgRNA sequence table, were provided as input. The output is a comma-separated value (CSV) file with one line per strain and well combination. The raw read counts were analyzed using DESEq2 ^70^ package in R, as previously mentioned^71^. Mutant proportions were determined as the percent ratio between the reads corresponding to each mutant and the total reads within a sample. Relative abundance was calculated as the percent ratio between each mutant proportion after and before pooled growth. Percent depletion was calculated as 100-relative abundance.

### Pth expression, purification and enzymatic activity analysis

Pth (K562_RS04080) was expressed from a pET22b(+) vector under the control of an IPTG-inducible promoter. First, *pth* gene was PCR amplified from wild-type *B. cenocepacia* K56-2 genome using Q5 polymerase with high GC buffer (NEB) and primers 3107 and 3108 (**Supplementary Table 10**) based on the following amplification condition: - 98°C for 30s, 30 cycles (98°C for 10s, 60.5°C for 20s and 72°C for 45s), and 72°C for 2mins. The 611bp amplicon and the pET22b(+) vector were double-digested with *NdeI* and *NotI*. Digested products were cleaned using Monarch^®^ PCR & DNA Cleanup Kit (NEB) and incubated with T4 DNA ligase (NEB) overnight at 16°C. The resulting plasmid, pPTH_BC_C2 was transformed into chemical-competent *E. coli* BL21(DE3) Gold cells, and ampicillin-resistant colonies were screened by colony PCR with primers 3008 and 3109.

For Pth purification, overnight cultures of *E. coli* cells harboring the plasmid were back diluted in 1:100 in LB (+100µg/mL ampicillin) and grown at 37°C with shaking until the OD600nm of 0.6 was reached. After that, isopropyl β-D-1-thiogalactopyranoside (IPTG) was added to a final concentration of 1mM to induce the protein expression and the culture was incubated at 20°C overnight. Cells were harvested via centrifugation, and the resulting cell pellets were resuspended in W/L buffer (50mM Tris-HCl pH 7.5, 500mM NaCl, 25mM imidazole) and stored at −80°C for at least 3 h. After thawing the pellet, 1mM phenylmethylsulfonyl fluoride (PMSF), 10mM MgCl_2_, and DNaseI was added to the suspension and incubated at room temperature for 5 mins. EmulsiFlex-C3 (Avestin) was used to lyse the cells. After centrifugation, the 6x His-tagged Pth_BC_ was purified from the soluble lysate via a nickel-NTA gravity column, followed by washing with 250ml of W/L buffer and elution with elution buffer (50mM Tris-HCl at pH 7.5, 500mM NaCl, and 500mM imidazole). The samples were passed through SnakeSkin (Thermo Scientific) 10K MWCO dialysis membrane overnight at 4°C for desalting and buffer exchange to remove imidazole.

Peptidyl-tRNA was synthesized and purified based on native chemical ligation (NCL) as described before^72^. All three compounds were resuspended to the desired concentration in 95:5 Ethanol: DMSO solution. 1nM Pth_BC_ was incubated in the reaction buffer (20mM HEPES-KOH, pH 7.2, 100mM NH_4_Cl, CH_3_COOK, 10mM MgCl_2_, and 1mM DTT) with formyl-Met-Thr-His-Ser-Met-Arg-Cys-tRNA^Cys^ (fMTHSMRC-tRNA^Cys^) in the presence (or absence) of the compounds. The hydrolysis reaction was quenched by mixing the sample with acidic gel loading buffer (Urea 7M, 100mM CH_3_COONa pH 5.2, 0.05% bromophenol blue, 0.05% xylene cyanole, 0.25% SDS, 10mM EDTA, 10% β-Mercaptoethanol) in 1:2 ratio. Before loading samples, the polyacrylamide gel (16×10 cm; 7M Urea, 12% acrylamide: bisacrylamide 19:1, 100mM CH_3_COONa pH 5.2) was pre-run in running buffer (100mM CH_3_COONa pH 5.2, 1mM EDTA) for an hour at 80V. The sample slots in the gel were washed right before loading the samples to aid in even loading and remove urea from the slots. The samples were run in the running buffer at 120V in a cold room (4°C) for 16 hours and stained with ethidium bromide (EtBr) solution in a staining tray for 5-10 minutes with shaking. The sensitivity of EtBr in acidic conditions is lower; therefore, before staining, the gels were washed several times in 1X TBE (2 minutes each wash). After staining, the gel was washed in 1X TBE for 5-10 minutes to reduce the background. The gel was visualized using the ChemiDoc MP imaging system (BioRad).

### Molecular docking

The 3D structure of *B. cenocepacia* K56-2 peptidyl-tRNA hydrolase (Pth_BC_) was generated with Alphafold^73^. The Pth_BC_ structure and the ligands were prepared for docking using Autodock Tools within Python prescription (PyRx) by adding polar hydrogen atoms and charges^74^. Subsequently, each prepared structure was converted into pdbqt format, which is compatible with AutoDock Vina. For docking, the Lamarckian genetic algorithm of Autodock Vina was employed to dock the ligands into the protein structure^75^. Following docking simulations, the model with the lowest energy was selected. The resultant complex was then aligned with the receptor using Pymol (Schrödinger, LLC) to visualize their spatial arrangement. 2D ligand-protein interactions were visualized with LigPlot+ v 2.2^76^.

### Data representation

All graphs, except for the clustering images in **Fig. 3C-D**, were created and generated using R (version 4.3.0). For the clustering graphs, analyses were performed using ‘sklearn.manifold’ and ‘sklearn.cluster’ packages in Python in an anaconda environment. The plots were visualized with ‘matplotlib.pyplot’ in Python. All correlations were calculated using the Pearson method unless otherwise noted. Schematics were created with Biorender web tool (https://www.biorender.com/). The band intensity of the cleaved and uncleaved peptidyl-tRNA was measured using ImageJ. Chemical structures in **Supplementary Fig. 11** were created with ChemDraw. Panels in the figures are compiled with Affinity publisher 2.

### Code and source data availability

All the raw read counts and source data for the figures and Python scripts (including the sgRNA design script) are available at https://github.com/zisanurrahman/CRISPRi_EGML_Main. The sequence of the closed K56-2 genome is under NCBI RefSeq accession: GCF_014357995.1.

## Acknowledgements

The authors thank Dr. Andrew M. Hogan for the insightful scientific discussions and for critically reading the manuscript. This work was supported by grants from the Canadian Institutes of Health Research (CIHR), Cystic Fibrosis Foundation, Cystic Fibrosis Canada to STC; the National Institutes of Health R01-GM132302 and R21-AI163466 to YSP, the National Science Foundation MCB-1907273 to YSP, and the Illinois State startup funds to YSP. ASMZR and JNN were supported by a University of Manitoba Graduate Fellowship (UMGF). AD was supported by a Mitacs Globalink Research Award. The funders had no role in study design, data collection and analysis, decision to publish or preparation of the manuscript.

## Declaration of generative AI and AI-assisted technologies in the writing process

During the preparation of this work the author(s) used Grammarly in order to improve grammar and clarity. After using this tool/service, the author(s) reviewed and edited the content as needed and take(s) full responsibility for the content of the publication.

## Author contributions

ASMZR – performed the majority of the experiments, analyzed data, and wrote the manuscript;

EAS – prepared peptidyl-tRNA, purified Pth_BC_, and performed Pth_BC_ activity assays;

JNN – contributed to the library preparation for CRISPRi-Seq and edited the manuscript;

AD – contributed to the hypersensitivity assay of the depleted mutants from pilot CRISPRi-Seq;

YSP – supervised the work, analyzed biochemical data, and reviewed the manuscript;

STC – conceived the idea, supervised the work, provided financial support, and edited the final version of the manuscript.

