## Supplementary information and supplementary figures for "Rationally Designed Pooled CRISPRi-Seq Uncovers an Inhibitor of Bacterial Peptidyl-tRNA Hydrolase"

***Supplementary Notes***

*Supplementary Note 1:* *Streptococcus pyogenes dCas9 (dCas9_Spy_) allows robust targeting in B. cenocepacia K56-2*

The PAM requirement of the CRISPR-Cas systems limits the targetable regions within a genome, prompting the exploration of orthologous (d)Cas9 proteins with different PAM preferences. While certain orthologous (d)Cas9 proteins can recognize multiple PAM sequences (**Supplementary Table 1**), the efficacy of most of these sequences remains insufficiently characterized. A handful of dCas9 orthologs have been characterized and applied for genetic manipulation. dCas9 orthologous proteins from *Streptococcus pasteurianus* (dCas9_Spa_) and *Streptococcus thermophilus* (dCas9_Sth1_) have been found to be efficient in GC-rich *Pseudomonas*^1^ and mycobacteria^2^, respectively. dCas9 orthologs preferentially recognize different PAM sequences within the DNA (**Supplementary Table 1**). Given the variable PAM requirement and efficacy of the dCas9 orthologs in diverse bacteria^1–6^, we aimed to select a suitable dCas9 ortholog that provides the greatest targeting flexibility to cover the *B. cenocepacia* K56-2 genome and allows efficient target knockdown. We performed a computational analysis to calculate the targetable regions (both total and 250bp upstream and 100bp downstream of CDS) in the *B. cenocepacia* K56-2 genome for the known (d)Cas9 orthologs (**Supplementary Table 1**). We shortlisted five candidate *Streptococcus* dCas9 orthologs that have been well-characterized for knockdown efficacy in bacteria: - *Streptococcus equinus* (dCas9_Seq_), *Streptococcus mutans* (dCas9_Smu_), *Streptococcus pasturianus* (dCas9_Spa_), *Streptococcus pyogenes* (dCas9_Spy_), and *Streptococcus thermophilus* CRISPR system I (dCas9_Sth1_). While dCas9_Seq_, and dCas9_Spy_ recognize short single PAM sequences, dCas9_Smu_, dCas9_Spa_, dCas9_Sth1_ recognize more than one PAM, albeit less preferentially^2,4,7^. Although dCas9_Smu_ can recognize NGG PAM, it preferred NAG with reduced activity in the presence of GC-rich NGG PAM^7^. Furthermore, variability in efficiency exists even within consensus PAM sequences. For instance, the consensus PAM sequence for dCas9_Sth1_ is NNAGAAW^8^ ("N" = any base and W = A or T). However, the efficiency varies across the characterized permissive PAM variants^2^. Additionally, the sgRNA design features for most of these orthogonal (d)Cas9 proteins are not extensively characterized. Considering that dCas9_Spy_ is the most extensively studied, has robust efficacy in our target organism, *B. cenocepacia*^9^, and possesses the simplest and GC-rich PAM preference ideal for GC-rich target organisms (i.e. *Burkholderia* has a GC content of ~67%^10^), we hypothesized that dCas9_Spy_ would allow the greatest coverage of the *B.* *cenocepacia* K56-2 essential genome. Indeed, *in silico* analysis of *B.* *cenocepacia* K56-2 genome against *Streptococcus* dCas9 orthologs revealed that dCas9_Spy_ has approximately 82, 5 and 2 times more targetable regions in the K56-2 genome compared to dCas9_Sth1_, dCas9_Spa_ and dCas9_Seq_, respectively (**Fig. 1B, Supplementary Table 1**). Moreover, we previously observed a strong knockdown efficiency of dCas9_Spy_ in *Burkholderia* species^9^. Given that dCas9_Spy_ provides a robust targeting coverage in *Burkholderia* without trading off the knockdown efficiency, we choose to use dCas9_Spy_ to construct a CRISPRi-based arrayed essential gene knockdown mutant library (EGML) in *Burkholderia cenocepacia* K56-2.

*Supplementary Note 2: Estimation of the percent depletion from clonal features*

The estimation of the percent depletion based on the random forest model provides insights into the relationship between clonal features and pooled depletion levels. Since we did not observe any apparent linear relationship between the clonal features and percent depletion, we performed feature engineering and converted the input clonal features into polynomial features to capture potential nonlinear relationships. Subsequently, a random forest regression model is fitted to the polynomial features, and predictions are made using the fitted model. By incorporating polynomial features, our model could better capture the complexities inherent in the relationship between clonal characteristics and depletion. The polynomial equation representing the relationship between the clonal features and pooled depletion is provided below:

$y$ *=c_0_+c_1_⋅* $x$*_1_+c_2_ ⋅* $x$*_2_+c_3_ ⋅* $x$*_1_^2^+c_4_ ⋅* $x$*_1_* $x$*_2_*

where,

$x$_1_ = µ_max(+Rhamnose)_, $x$_2_ = lag time, *c_0_* = intercept of the hyperplane, *c_1_* = coefficient for $x$_1_, *c_2_* = coefficient for $x$_2_, *c_3_* = coefficient for $x$_1_^2^, *c_4_* = coefficient for $x$_1_ *⋅* $x$_2_

*Supplementary Note 3: Kernel PCA*

PCA aims to find a lower-dimensional representation of high-dimensional data while preserving the variance in the data as much as possible. However, PCA assumes that the data is linearly separable, which may not always be true for complex datasets. Kernel PCA addresses this limitation by employing a kernel function to implicitly map the original data into a higher-dimensional feature space where it may be linearly separable. In this feature space, PCA is then applied to extract principal components.

**Supplementary Figures**

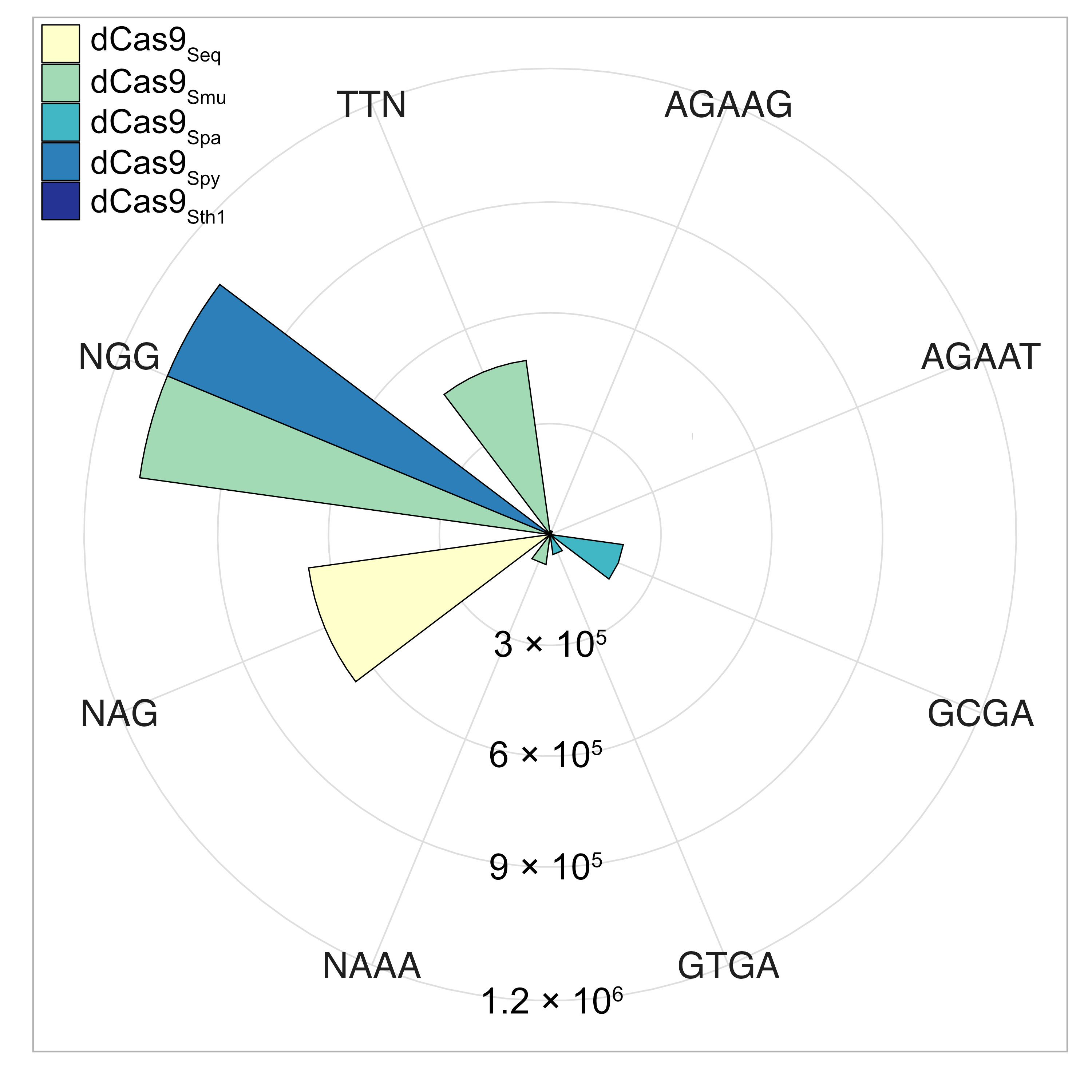

**Supplementary Fig. 1:** Number of possible target sites in *B. cenocepacia* K56-2 genome using orthologous dCas9 proteins.

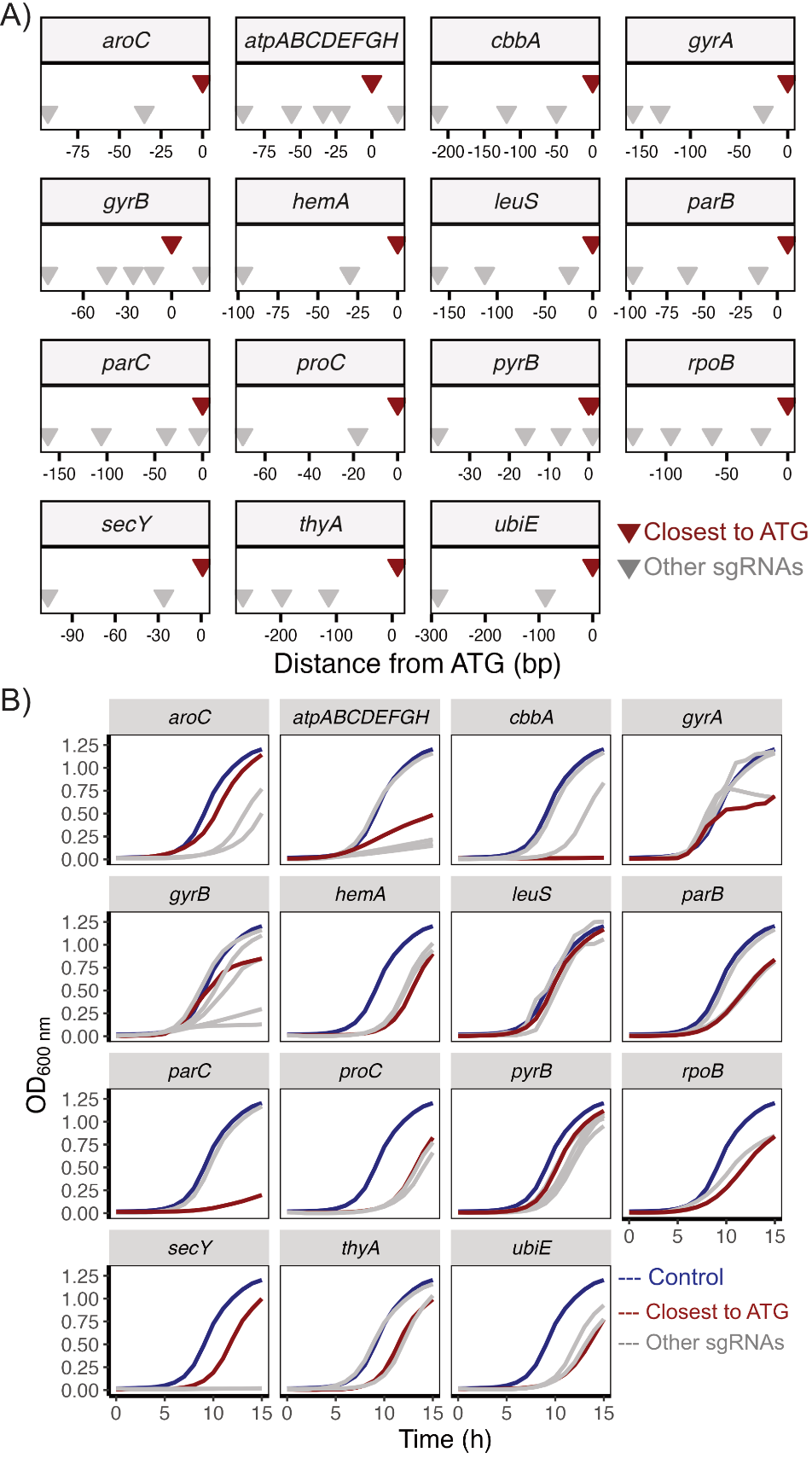

**Supplementary Fig. 2: Pilot study to deduce a genomic target region for efficient gene repression.** A) Relative sgRNA location targeting different regions of 15 selected genes. sgRNAs were designed to target the non-template strand. B) Growth curves of the CRISPRi mutants created with the sgRNAs shown in panel A. Overnight cultures of the mutants were OD_600nm_ adjusted to 0.01 and grown with and without 1% rhamnose in a plate reader, and reading was taken at one-hour interval. Results are the mean of 3 biological replicates, and error bars represent the mean ± SD.

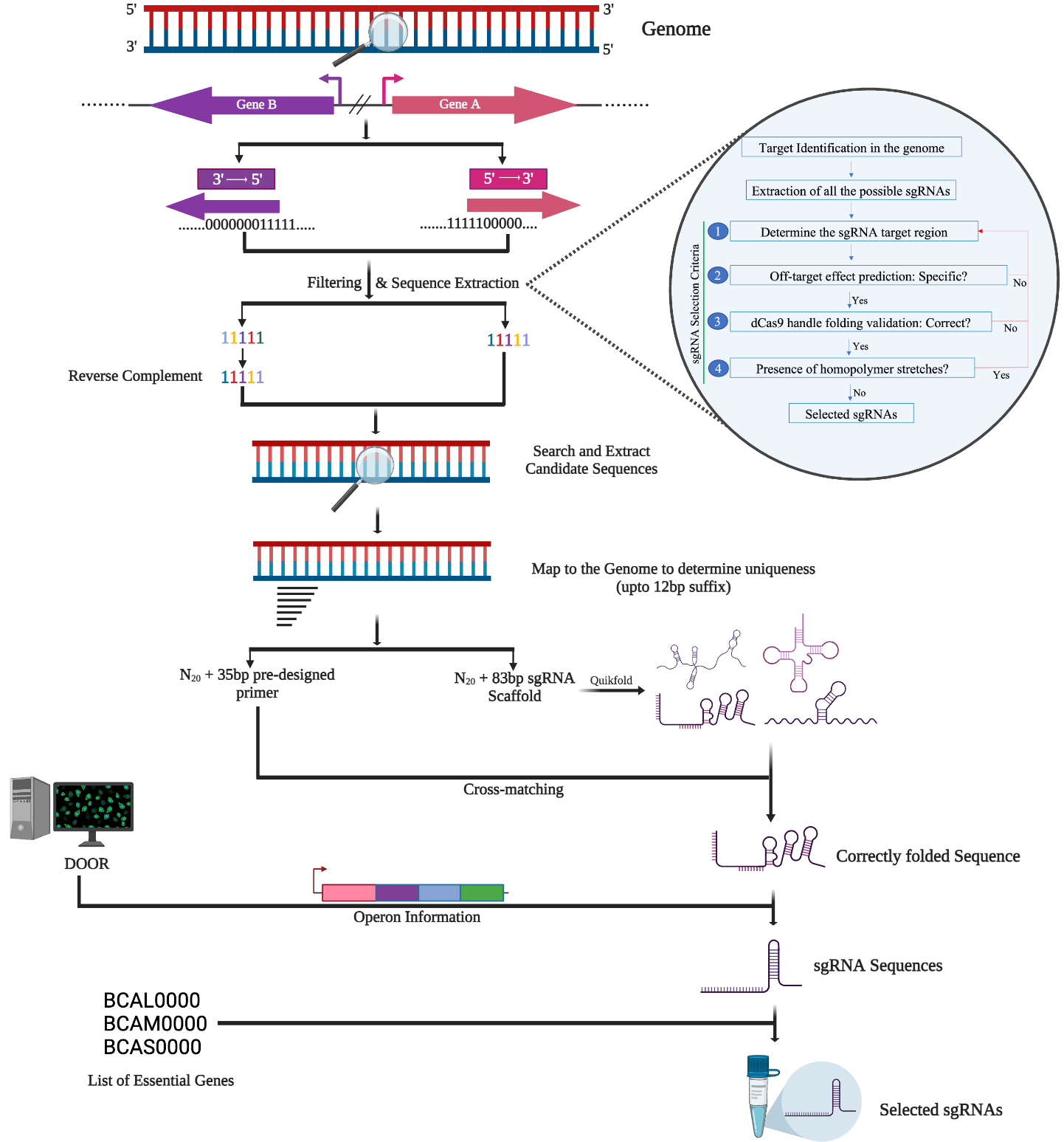

**Supplementary Fig. 3:** **Custom Python script to computationally design sgRNAs targeting the essential genome of *B. cenocepacia* K56-2.** The script goes through a series of steps, and the output is ready-to-order primers to clone the sgRNAs. At first, the script splits the genome into two groups, each facing the opposite direction, and extracts the sequence based on a provided query range. For example, if the script is provided a query range of -100 to +50 for a given gene, the script will extract a total of 150 bp, 100 bp upstream and 50 bp downstream of the translation start site (ATG). The script will then look for the possible sgRNAs based on the availability of the PAM sites and extract the candidate sgRNAs. The candidate 20bp sgRNAs are then mapped to the genome to determine the specificity by paring one base at a time from the 3’ end up to 12bp and will select the one sgRNA per gene that does not have off-target binding sites, free of homopolymer stretches (e.g. more than 5) and closest to the ATG site. The script is customizable and can be modified to target the whole genome or only the essential genome, alter the query range, and provide more than one sgRNA per gene, if necessary. The provided sgRNAs are 20bp long, which are then concatenated with the sgRNA scaffold sequence to predict the secondary structure folding of the hairpin required for dCas9 binding using Quikfold algorithm from the UNAFold package. Successful sgRNAs are then concatenated with the pre-designed forward primer as a 5’ extension and used for cloning the sgRNAs.

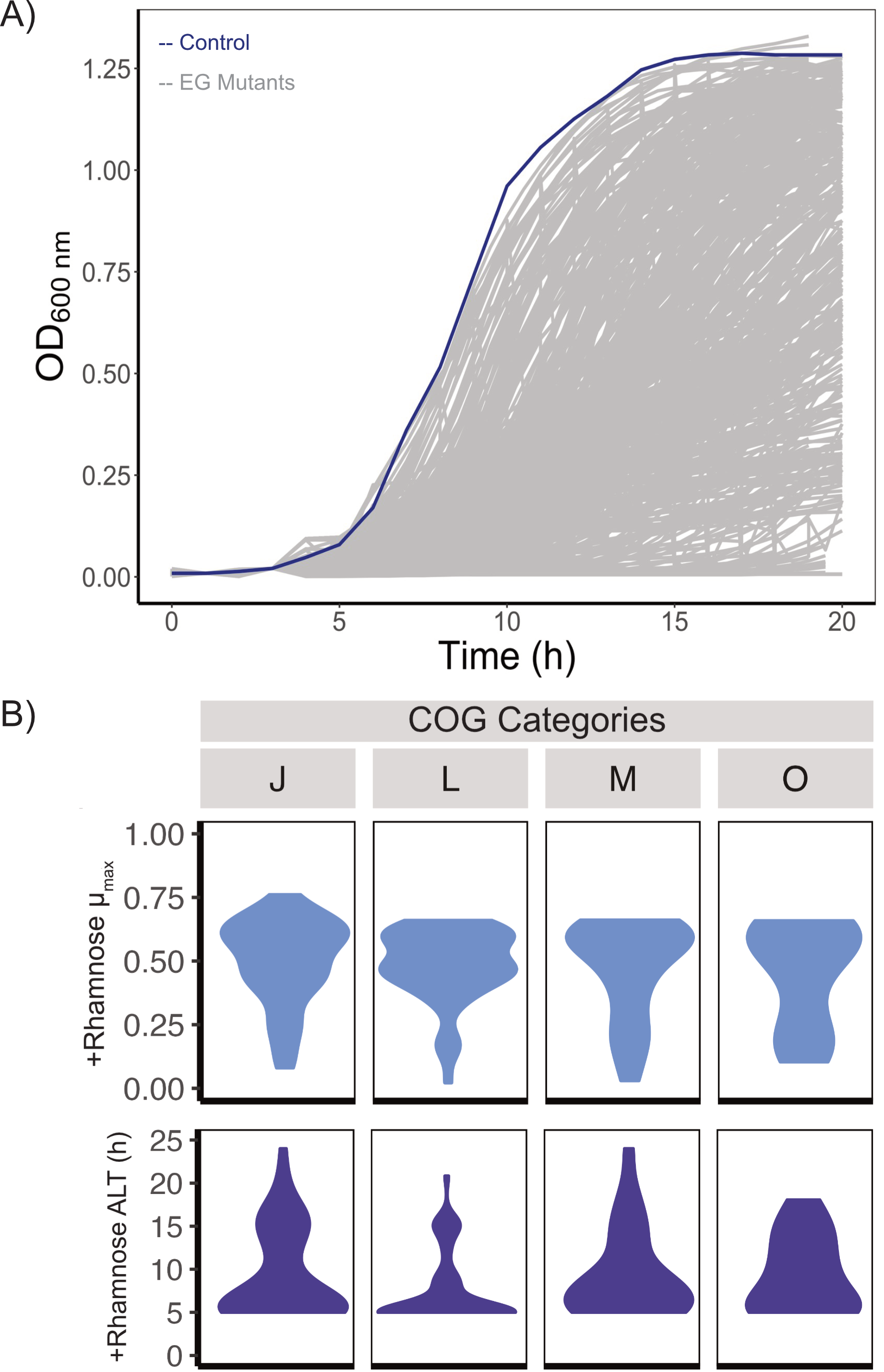

**Supplementary Fig. 4: Growth characterization of *B. cenocepacia* K56-2 CRISPRi essential gene mutants**. A) Microplate reader growth curves of the knockdown strains. The lines are the mean of at least three independent biological replicates. The blue line indicates the pgRNA-nontarget control (n = 20), and each grey line indicates individual essential gene mutants. B) Clonal maximum growth rate (µ_max_) and apparent lag time (ALT) of the mutants (in inducing condition) belonging to COG categories related to translation (COG J and O), membrane biogenesis (M) and DNA replication and repair (L). ALT denotes ‘apparent lag time’. µ_max_ and apparent lag time were calculated with the ‘growthrates’^11^ package in R.

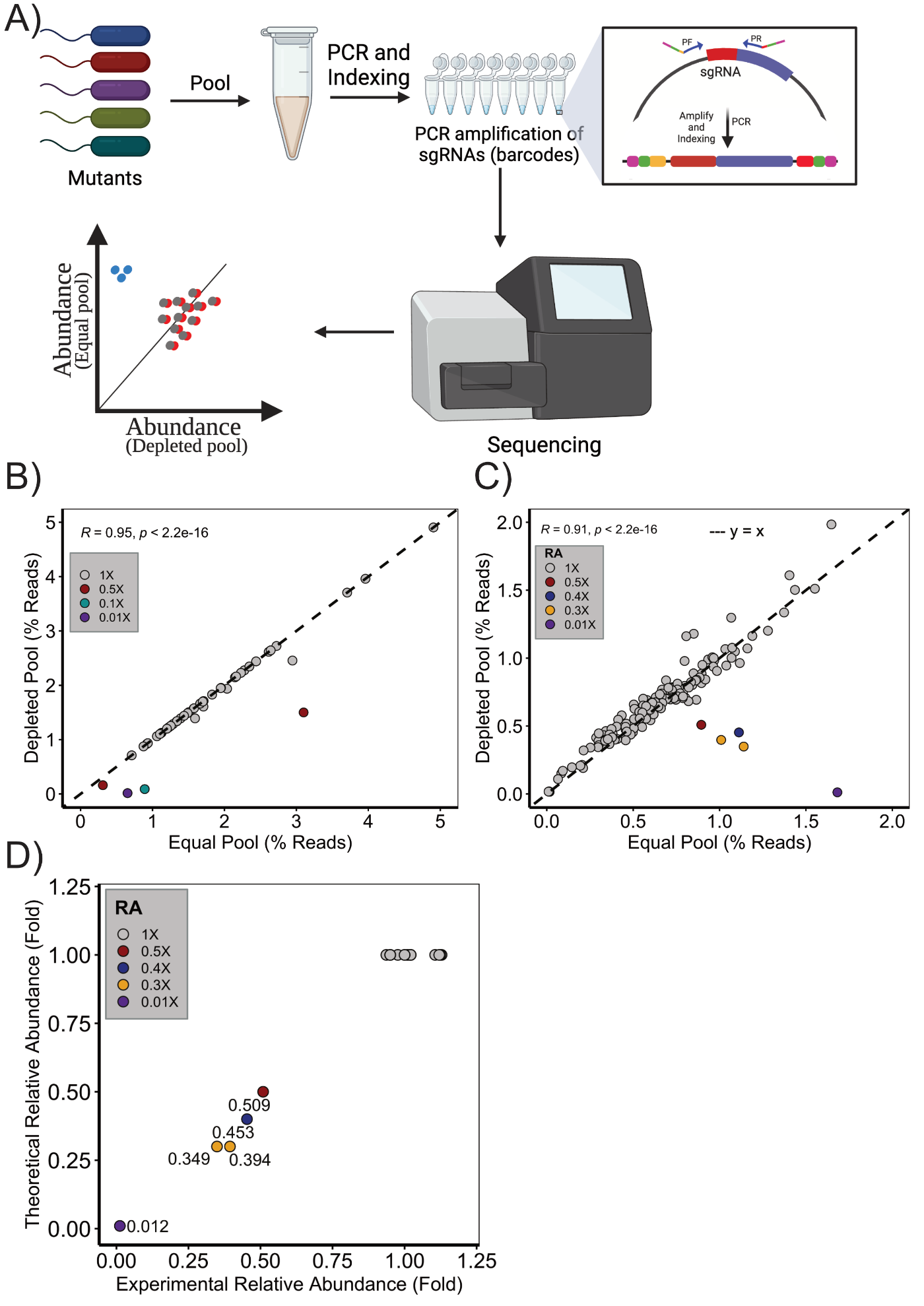

**Supplementary Fig. 5: CRISPRi-Seq approach to detect the change in relative abundance of specific mutants.** A) Overview of the CRISPRi-Seq approach to detect changes in the relative abundance of the mutants by Illumina sequencing of sgRNA-encoding DNA barcode counts. B-C) Relative abundance (compared to the equal pool) of the mutants from the pool of 50 mutants (B) and 150 mutants (C), where four and six mutants were artificially depleted, respectively. D) The normalized (experimental) relative abundance from CRISPRi-Seq is consistent with the theoretical relative abundance for the mutants. The result is the mean of two independent biological replicates. The correlation coefficient (R) was calculated based on the Pearson method.

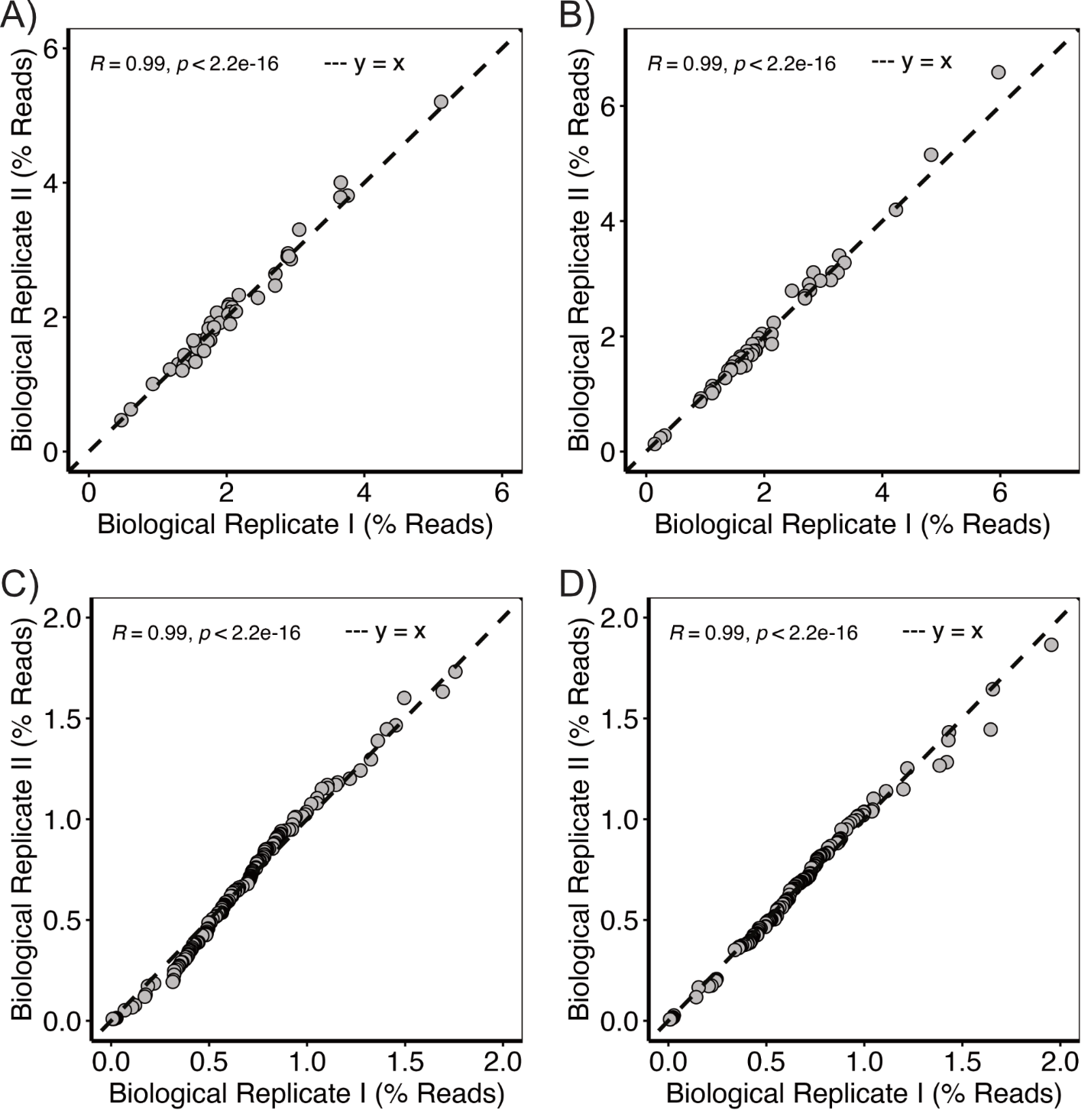

**Supplementary Fig. 6:** **Correlation of relative abundance across biological replicates**. The percent abundance of each mutant in the equal pools (A and C) and depleted pools (B and D) of 50 and 150 mutants, respectively. The relative abundance was consistent across the biological replicates, showing that each mutant was reproducibly amplified by PCR and detected. The correlation coefficient (R) was calculated based on the Pearson method.

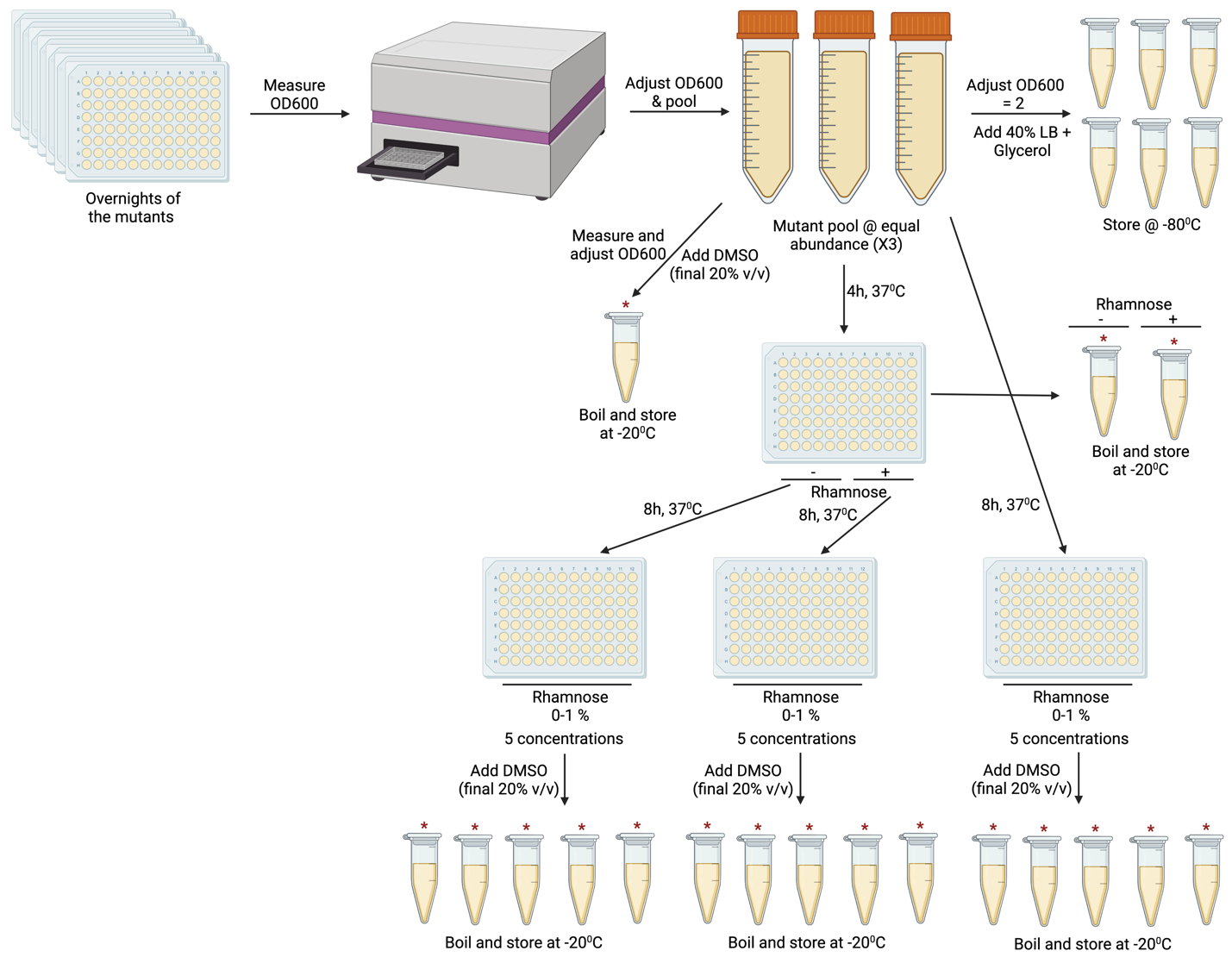

**Supplementary Fig. 7:** **Workflow to determine the suitable sensitizing rhamnose concentrations from ‘equal EGML’ to bin the mutants.** Mutants were pooled and grown in five different rhamnose concentrations ranging from 0 to 1%, followed by Illumina sequencing of the pools. The proportions of reads for the mutants in a pool were used as a proxy to determine the depletion of the mutants from a specific pool. Starred tubes represent the sequenced samples.

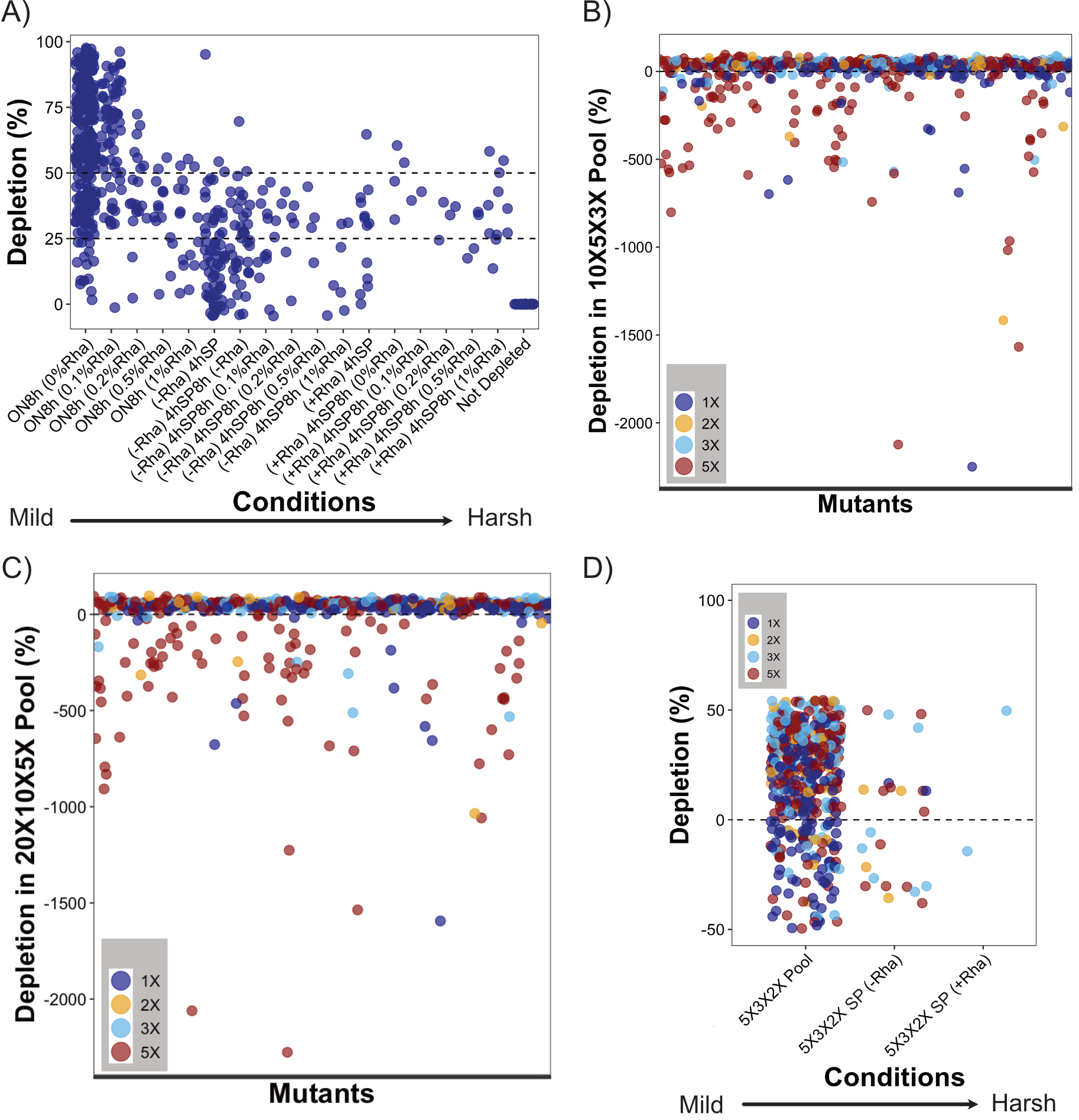

**Supplementary Fig. 8: Characterization of the mutants in pooled competitive growth conditions to create an optimized pool.** A) Depletion of mutants from 1X (equal) pool in response to rhamnose. Clonal overnight cultures of the mutants were binned and grown for 8h (additional 4h in case of sub-culture) at different rhamnose concentrations ranging from 0% to 1%. Each dot represents a unique mutant. The mildest condition at which the mutants first depleted are plotted. The vertical dashed grey lines represent the 25-50% depletion. The abbreviation "ON" denotes overnight, "8h" signifies growth after pooling, and the percentage of rhamnose (in parentheses) indicates the concentration at which the pools were grown for 8 hours. SP denotes ‘sub-cultured pool’ where the pool was grown in the absence (-Rha) or presence (+Rha) for an additional 4h before the 8h growth. (B-C) Depletion of the ‘sick’ and ‘highly sick’ mutants (from 1X pool) in the spiked 10X5X3X (B) and 20X10X5X (C) pools. The mutant relative abundance (RA) in the pools was calculated based on the proportions of reads for the mutants in a pool and then compared to a control pool that had not been grown after pooling. D) Depletion of mutants from the 5X3X2X pool. The 5X3X2X pool was grown for 8h (Pool) or 8 h after previously pooled subculturing for 4h (SP) in the absence (-Rha) or presence (+Rha) of 0.1% rhamnose. Each dot represents a unique mutant. The mildest condition at which the mutants first depleted are plotted. Negative depletion indicates enrichment of the mutants compared to the control. Results are the mean of at least two independent biological replicates.

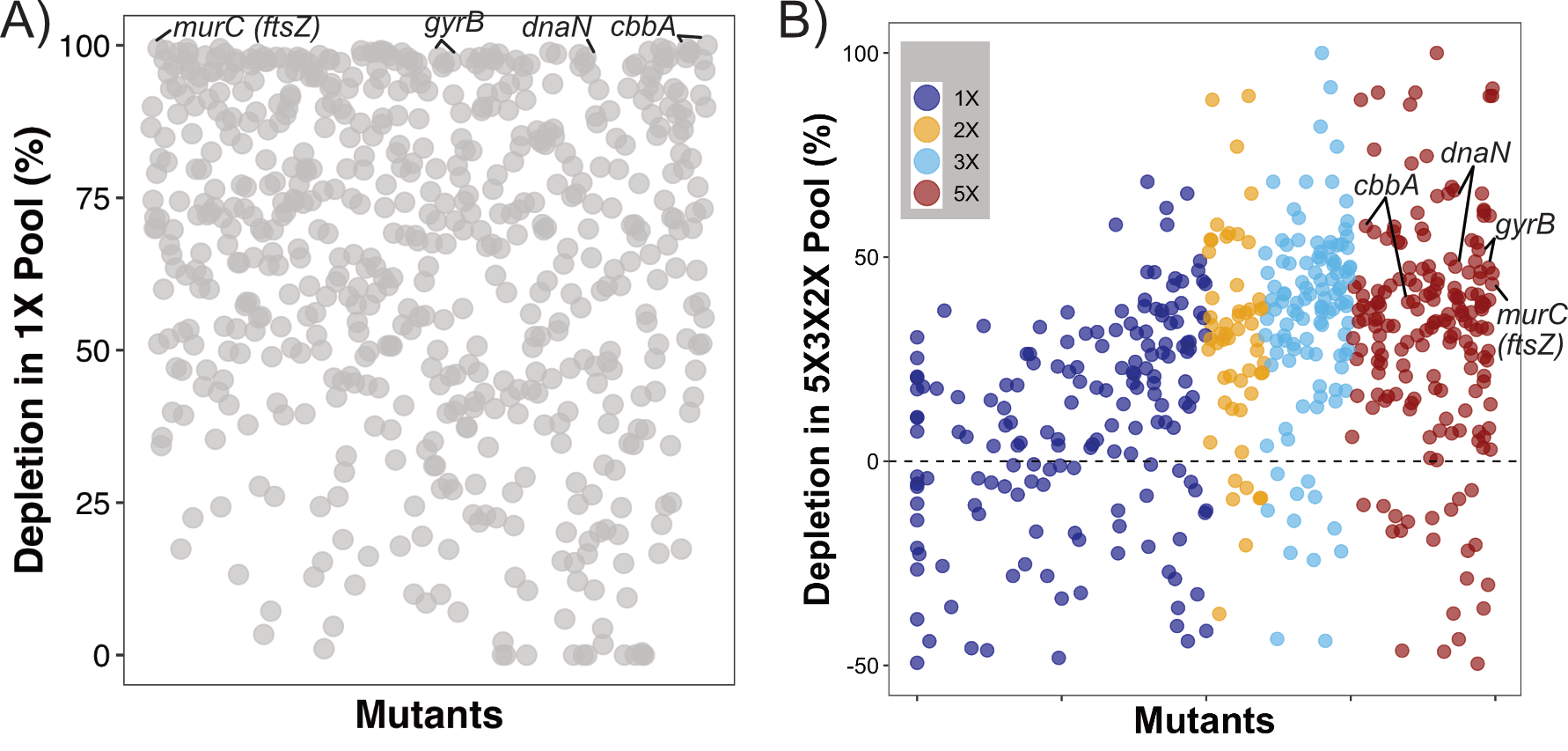

**Supplementary Fig. 9: Mutant depletion from a 1X pool (A) and rationally spiked 5X3X2X pool (B).** Each dot represents a unique mutant. The relative abundance (RA) of mutants in the pools was determined by calculating the proportions of reads for each mutant in a pool and comparing them to a control pool that had not been grown after pooling. Negative depletion indicates enrichment of the mutants compared to the control. Results are the mean of at least two independent biological replicates.

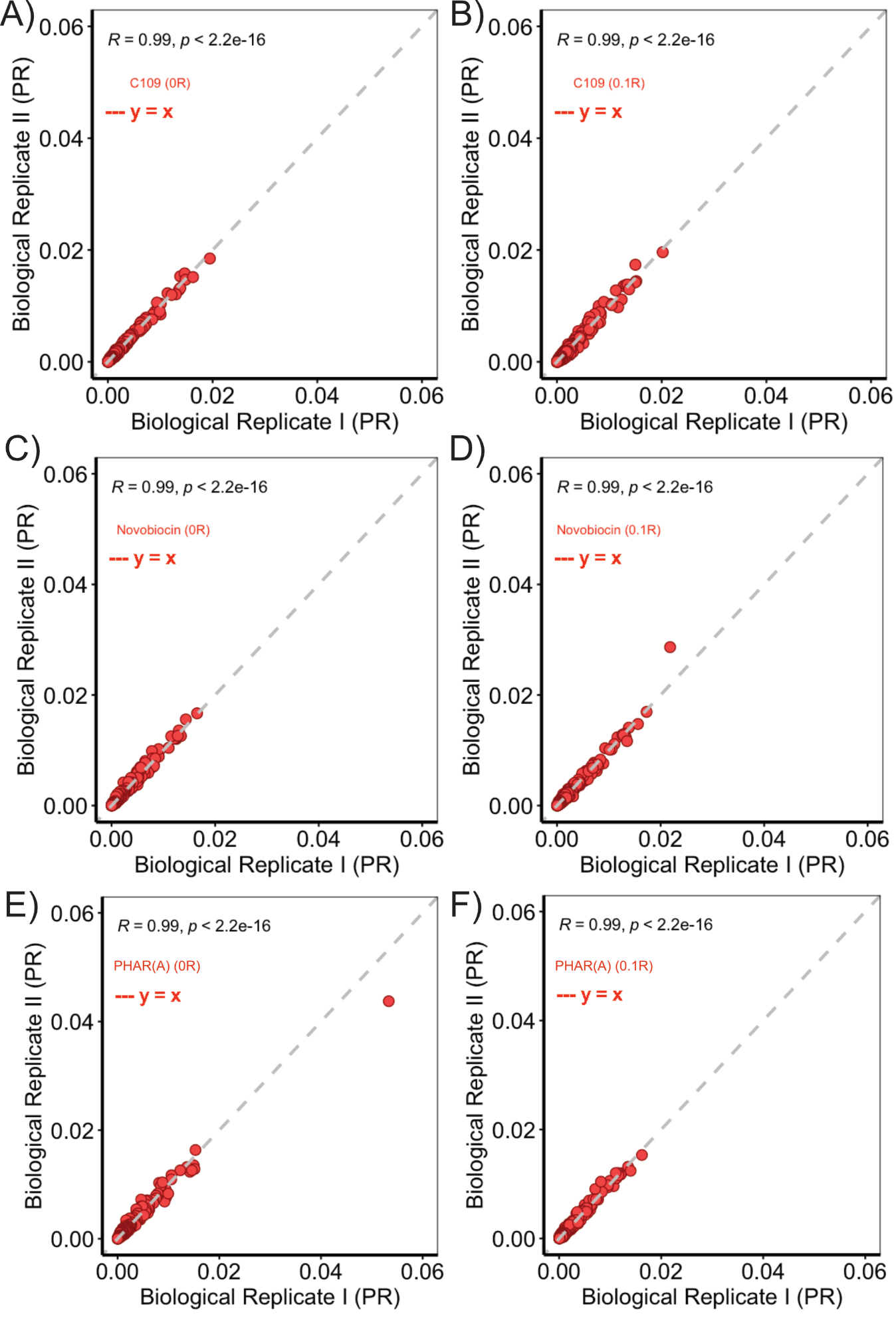

**Supplementary Fig. 10: Correlation of proportion of reads (PR) across biological replicates**. The proportion reads of each mutant between the two biological replicates in the optimized EGML (5X3X2X) pool after treatment with C109 (A-B), novobiocin (C-D) and PHAR(A) (E-F). EGML exposure to antibiotics without (B and D) and with (C and E) of 0.1% rhamnose. EGML exposure to antibiotics without (A, C, and E) and with (B, D, and F) of 0.1% rhamnose. The correlation coefficient (R) was calculated based on the Pearson method.

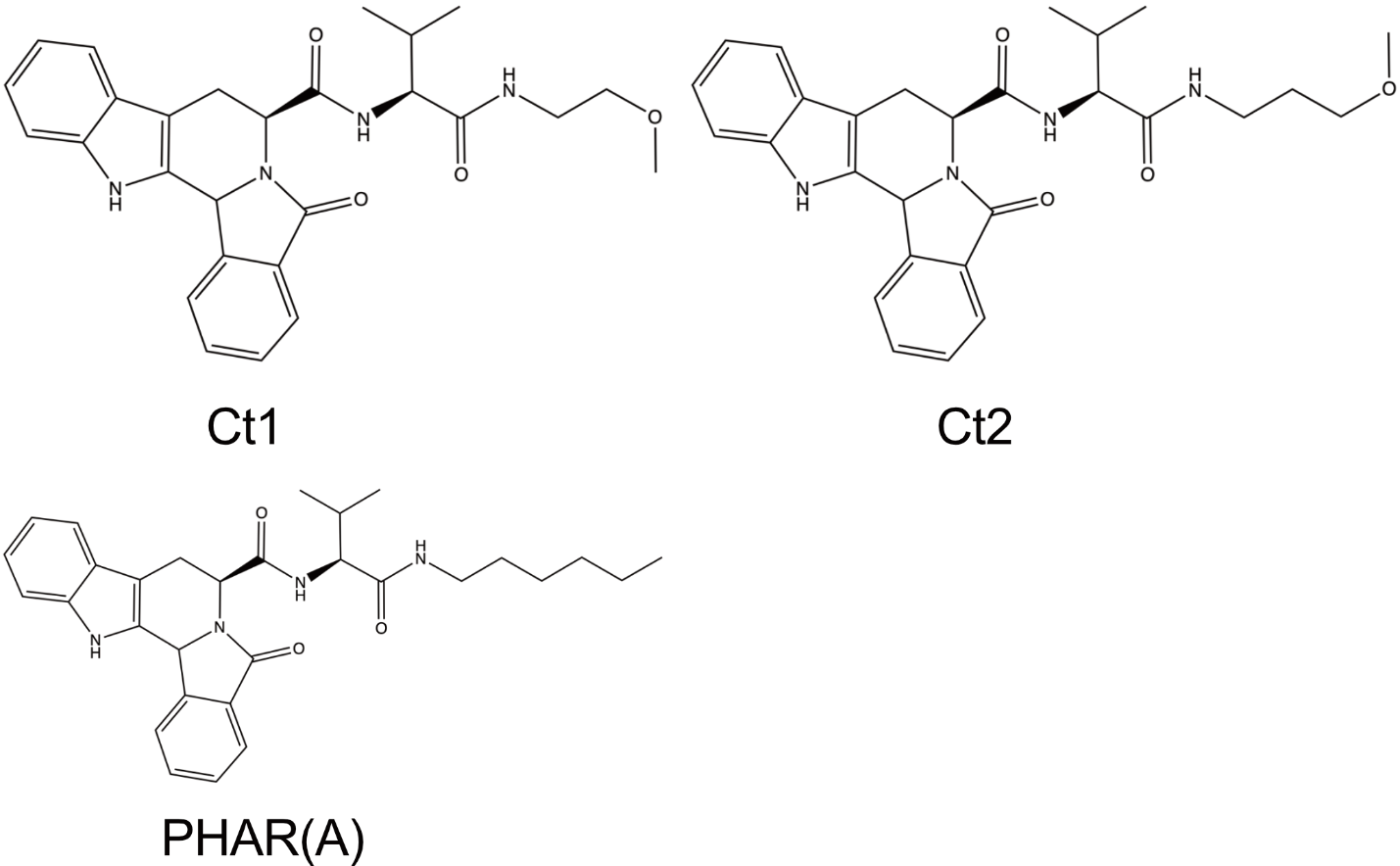

**Supplementary Fig. 11:** Chemical structure of PHAR(A) and the control (Ct) compounds.

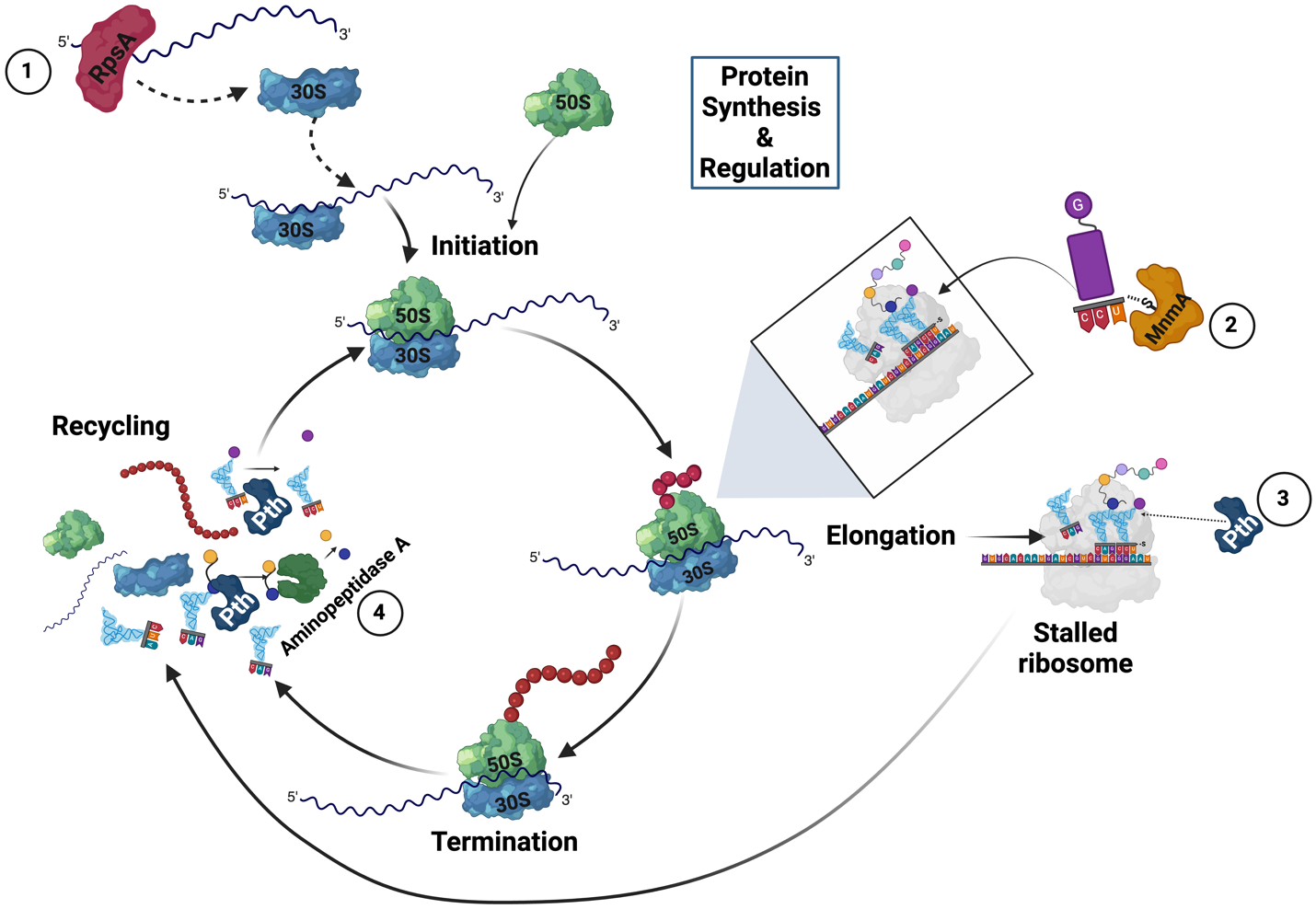

**Supplementary Fig. 12:** **Inhibition of peptidyl-tRNA hydrolase by PHAR(A) disrupts the overall translation process.** RpsA (1), MnmA (2), Pth (3), and aminopeptidase A (4) are involved in different stages of protein synthesis and recycling.

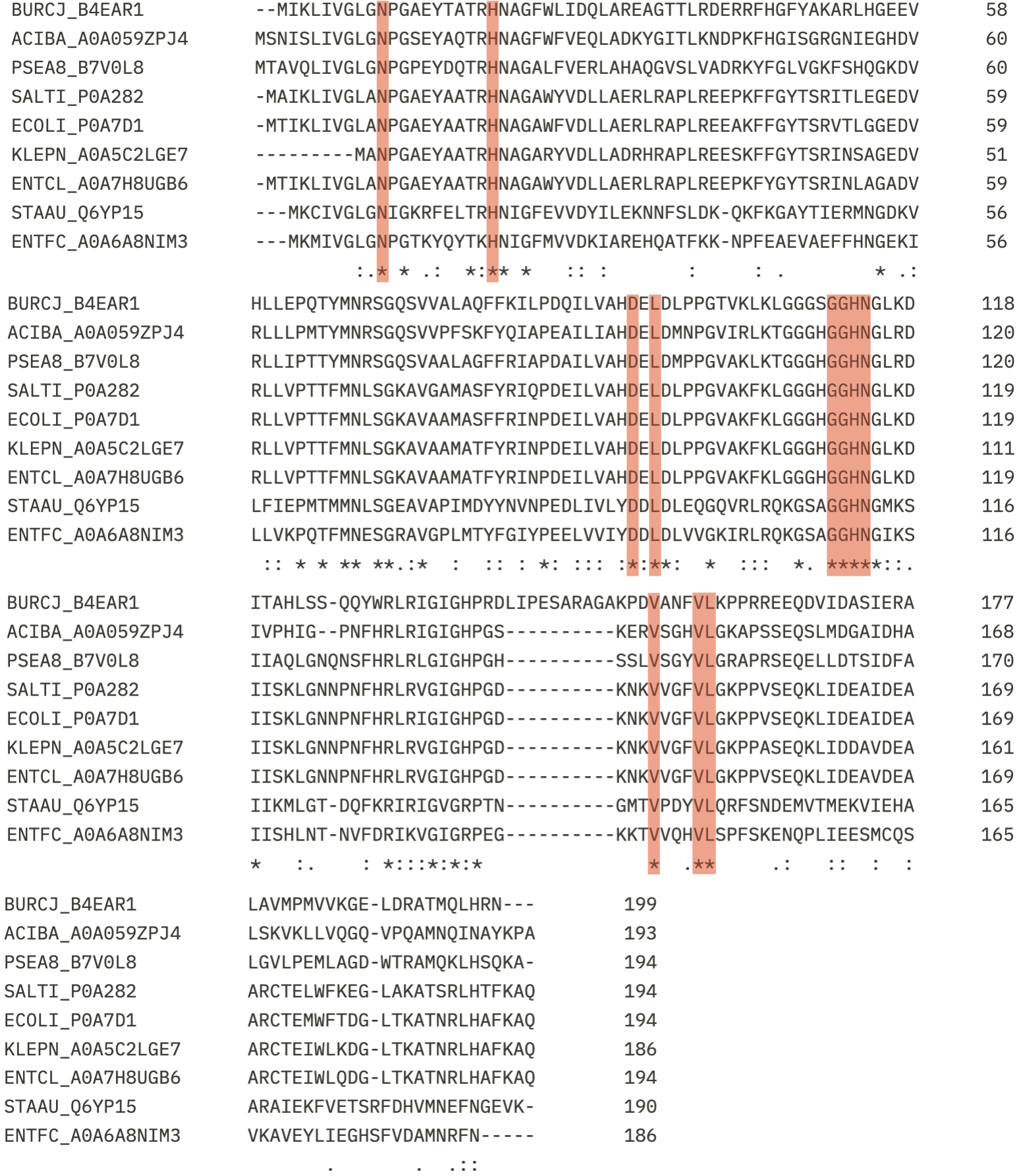

**Supplementary Fig. 13:** **Multiple sequence alignment of peptidyl-tRNA hydrolase (Pth) from diverse bacterial species.** Protein sequences were collected from UniProt database (<https://www.uniprot.org/>) and aligned using ClustalΩ within EMBL-EBI (<https://www.ebi.ac.uk/jdispatcher/msa/clustalo>). Amino acid residues involved in the interactions with PHAR(A) and/or its inactive analogs are highlighted. Pth sequence from *Burkholderia cenocepacia* (BURCJ), *Acinetobacter baumannii* (ACIBA), *Pseudomonas aeruginosa* (PSEA8), *Salmonella typhi* (SALTI), *Escherichia coli* (ECOLI), *Klebsiella pneumoniae* (KLEPN), *Enterobacter cloacae* (ENTCL), *Staphylococcus aureus* (STAAU), *Enterococcus faecium* (ENTFC).

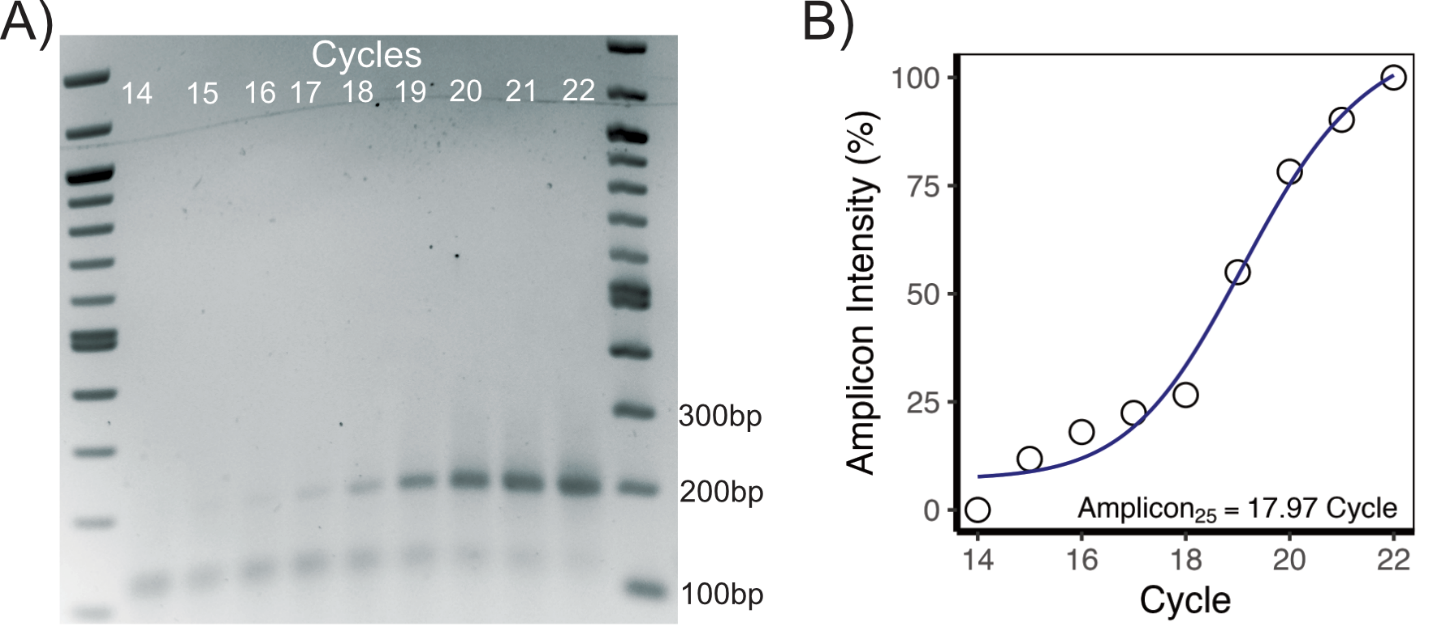

**Supplementary Fig. 14:** **Semi-quantitative PCR to determine the cycle number corresponding to ~25% maximum product (Amplicon_25_)**. A) Gel image of the PCR products after different PCR cycles. Samples were electrophoretically separated in 2.5% agarose gel, run for 29 mins at 190V and stained with ethidium bromide. B) Amplicon_25_ was determined based on the amplicon band intensity at various PCR cycles. The band intensity was measured using ImageJ ^12^ and the Amplicon_25_ was calculated using the “DRC” ^13^ package in R.

**Supplementary Methods**

*Antibiotics and bioactive compounds formulations*

C109 was synthesized following the procedure outlined in Scoffone et al. (2015)^14^. All the antibiotics and the bioactive compounds with unknown mechanisms of action used in this study were dissolved in dimethyl sulfoxide (DMSO) at various stock concentrations. The stock and the testing concentrations are provided in **Supplementary Table 7**. 1M Isopropyl ß-D-1-thiogalactopyranoside (IPTG) and 20% L-rhamnose (Sigma) solutions were prepared in water.

*Cloning the sgRNAs and construction of the CRISPRi EGML in a high-throughput manner*

The 20bp sgRNA sequences were cloned into the sgRNA expressing vector, pgRNA, by inverse PCR in a 96-well PCR plate (Sarstedt). The linear PCR products include the plasmid components and the 20bp sgRNA sequences. 0.5µl of the PCR products were added to a 96-well PCR plate pre-filled with 4.5µl quick ligation mastermix (0.5μl *DpnI*, 0.5μl T4 polynucleotide kinase, and 0.5μL T4 ligase with quick ligation buffer (NEB)) and incubated for 30mins at 37^0^C. 2µl of the ligation products were promptly added to *E. coli* DH5α competent cells and transformed by heat-shocking at 42^0^C for 45s. *E. coli* DH5α competent cells were prepared in advance and stored in 96-well PCR plates at -80^0^C and thawed on ice before use. The transformation mixtures were pinned into 9cm × 9cm square petri dishes (Sarstedt) in an 8 × 6 matrix. The transformants were selected in trimethoprim (50µg/mL) containing LB plates and screened using primers 1409 and 848, which bind upstream and downstream of the sgRNA sequences. The *E. coli* DH5α cells harbouring the sgRNA plasmid, *E. coli* MM294/pRK2013 helper strain and the *B. cenocepacia* K56-2 mutant expressing dCas9_Spy_ (K56-2::*dCas9_Spy_*) were mated on 9cm × 9cm square petri dishes (Sarstedt) with LB in an 8 × 6 matrix. The transconjugants were selected in trimethoprim (100µg/mL) containing LB plates and confirmed by colony PCR with primers 1409 and 848.
